# Unifying coexistence theory for ecological communities and pathogen strain competition

**DOI:** 10.64898/2026.05.27.728210

**Authors:** Sang Woo Park, Jonathan Levine, Bryan T. Grenfell

## Abstract

Predicting the outcome of species or pathogen strain competition is a fundamental aim in both community ecology and infectious disease dynamics. Recent work revealed major challenges in predicting strain co-circulation from ecological coexistence theory due to overcompensatory competition among pathogens for susceptible resources, which can prevent the re-invasion of other competing strains. This resource overcompensation is ubiquitous across host-pathogen systems, but not apparent in simple Lotka-Volterra competition system, highlighting fundamental differences between pathogen strain and species competition. To address this gap, we begin by deriving classical models of pathogen strain and species competition from a resource-consumer model. This generalization illustrates that the relative time scale between resource and consumer dynamics limits the degree of resource overcompensation and therefore dictates the outcome of stochastic competition. Moreover, by introducing a mathematical framework for quantifying pairwise and higher-order terms from general competition systems, we show that a simple, ecological competition model can accurately predict the equilibrium dynamics of strain competition. A case study of rotavirus strain competition reveals that the ability to predict the outcome of strain competition from ecological theory depends on the underlying cross immunity structure. This work synthesizes coexistence theory across two fields by providing a unifying framework for predicting the outcome of complex ecological competition.

## Introduction

Understanding what factors determine species or strain coexistence is a major question in both community ecology (Hutchinson, 1961; Chesson, 2000; Johnson et al., 2022) and infectious disease dynamics (Cobey and Lipsitch, 2012; Lloyd-Smith, 2013; Rice et al., 2021; Sieben et al., 2022; Park et al., 2024). However, fundamental differences in mechanisms underlying species competition and pathogen strain circulations have led to a development of disparate mathematical theory and modeling tools across two fields. For example, the Lotka-Volterra competition model represents a core approximation of complex ecological communities and provides a foundational basis for understanding species competition (Volterra, 1927; Lotka, 1932; May and Leonard, 1975; Chesson, 2000; Levine et al., 2017; Gibbs et al., 2022). In contrast, characterizing pathogen strain competition often requires detailed mechanistic models that are finely tuned for each host-pathogen system (Gog and Grenfell, 2002; Ferguson et al., 2003; Koelle et al., 2006; Pitzer et al., 2011; Bhattacharyya et al., 2015; Howerton et al., 2025). A unifying framework for explaining the differences and similarities in the dynamics of species and strain competition remains elusive.

One of the main features of pathogen strain competition, which appears to be absent in the Lotka-Volterra competition model, is resource overcompensation for susceptible hosts. Specifically, when an outbreak occurs, an invading pathogen causes a major susceptible depletion that continues even after the epidemic peak, causing the amount susceptible population to reduce below the invasion threshold (Restif and Grenfell, 2006, 2007; Ballesteros et al., 2009). This resource overcompensation can therefore cause an extinction of both invading and resident strains and prevent coexistence for mutually invading strains (Park et al., 2024). This phenomenon provides a foundation for understanding why some pathogen strains, such as RSV A vs B, can co-circulate, while influenza strains and SARS-CoV-2 variants exhibit strain replacement despite their ability to mutually invade one another (Park et al., 2024). This observation appears to contradict the classical ecological theory that mutually invading species should coexist (Chesson, 2000). So what mechanisms drive the resource overcompensation and why it is so prevalent in host-pathogen systems, but not apparent in the Lotka-Volterra competition model? And can simple ecological theory still provide useful predictions for multistrain pathogen systems?

To answer these questions, we present a unifying framework for linking pathogen strain competition to ecological species competition. We begin by showing that a classical model for multi-strain dynamics can be obtained from a general consumer-resource model. The Lotka-Volterra competition model can be then derived from the multi-strain model by assuming equilibrium resource dynamics and applying the first-order Taylor approximation around equilibrium. In doing so, we illustrate that the relative speed between resource and consumer dynamics affects resource overcompensation and determines the outcome of competition. This derivation further illustrates discrepancies in predictions about equilibrium densities between the multistrain model and the Lotka-Volterra competition model based on the Taylor approximation around equilibrium. Moreover, deriving a Lotka-Volterra competition model from more complex multistrain systems may not be always feasible, necessitating a more general approach for linking realistic competition systems of pathogen strain dynamics with the Lotka-Volterra competition model. To address these gaps, we develop a general framework for quantifying pairwise and higher-order interactions from general competition models and apply it to our multistrain model, revealing how higher-order interactions emerge from nonlinear host-pathogen strain interactions and determine equilibrium dynamics. Finally, to understand how the nature of competition shapes the utility of Lotka-Volterra model in predicting equilibrium dynamics of more complex systems, we apply the resulting framework to rotavirus strain dynamics.

### Mathematical theory

To understand the differences between species and strain competition, we begin with a general consumer-resource model (Letten and Stouffer, 2019). Specifically, we consider *n* consumers, *N*_*i*_, competing for *n* resources, *R*_*j*_ (Figure 1A; Supplementary Text S1):

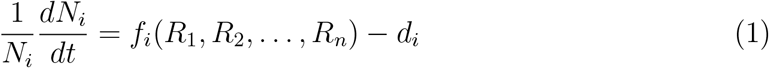

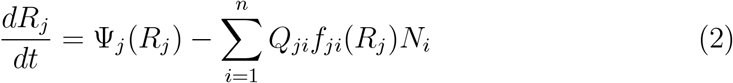

where *f*_*i*_ represents the per capita growth rate of consumer *i*; *d*_*i*_ represents the per capita death rate of the consumer *i*; Ψ_*j*_ represents the replenishment rate of the resource *i*; and *Q*_*ji*_ represents the amount of resource *j* that is used by consumer *i*.

**Figure 1:**
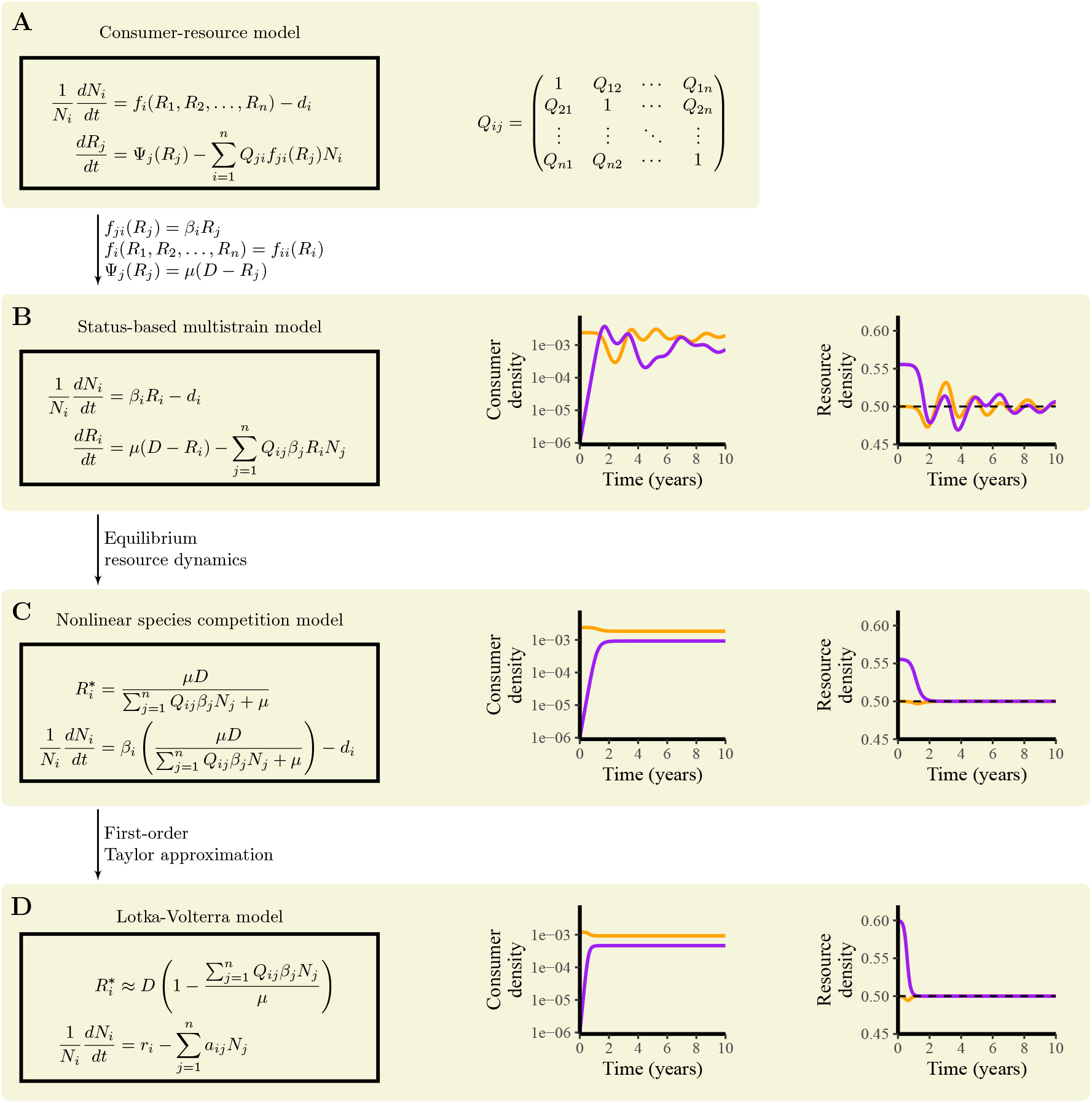
Schematic diagram illustrating different model formulations and the resulting competition dynamics. (A) A general consumer resource model describing species competition. (B) A classic multistrain model is a special case of a consumer resource model that assumes (1) linear resource consumption, (2) strain-specific resource limitation, and (3) linear resource replenishment. (C) Assuming equilibrium resource dynamics removes cycles in strain competition. (D) Applying the first-order Taylor expansion yields a simple Lotka-Volterra model. See Supplementary Figure S1 for a side-by-side comparison of equilibrium densities across three models, illustrating underestimation from the Lotka-Volterra model.

To derive a multistrain host-pathogen model from the resource-consumer model, we make three simplifying assumptions: (1) per capita growth rate of consumer *i* is a linear function of the resource, *f*_*ji*_(*R*_*j*_) = *β*_*i*_*R*_*j*_; (2) resource *i* limits the growth of consumer *i* such that the growth of consumer *i* only depends on resource *i, f*_*i*_(*R*_1_, *R*_2_, …, *R*_*n*_) = *f*_*ii*_(*R*_*i*_); and (3) resource becomes replenished at a fixed rate, Ψ_*j*_(*R*_*j*_) = *µ*(*D* − *R*_*j*_). Notably, this constant resource replenishment assumption has been commonly used for modeling abiotic resources and chemostat competition (Tilman, 1977, 1980; Hsu and Waltman, 1998) as well as for modeling susceptible human populations (Anderson and May, 1979; May and Anderson, 1979)—in both scenarios, the resource term gets replenished at a constant rate, independent of the consumer dynamics. Under these assumptions, the consumer-resource model simplifies to (Figure 1B):

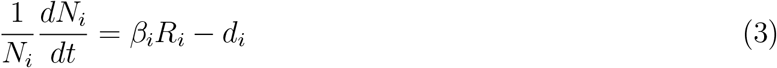

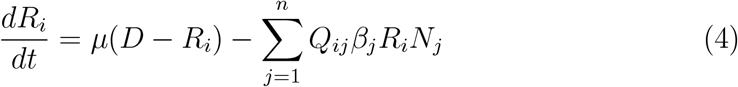

where *R*_*i*_ now represents the density of susceptible hosts (resource); *N*_*i*_ represents the density of infected hosts (consumer); *Q*_*ij*_ represents the degree of cross immunity that infection by strain *j* confers on infections by strain *i*; *β*_*i*_ represents the transmission rate of strain *i*; *d*_*i*_ represents the mean clearance rate of infection by strain *i*; and *D* represents the total population size. This model formulation corresponds to a status-based representation of a multistrain system (Gog and Grenfell, 2002) and is also broadly similar to a consumer-resource model formulation used previously in the context of studying higher-order interactions (Letten and Stouffer, 2019). As we discuss in Supplementary Text S2, the parameter *µ* captures the effective rate of susceptible replenishment, combining both birth rates and waning immunity rates.

If we now take this model and further assume that the resource dynamics occur at a much faster rate than the consumer dynamics such that the resource is always at equilibrium, we obtain a nonlinear species-competition model (Figure 1C):

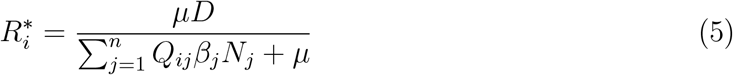

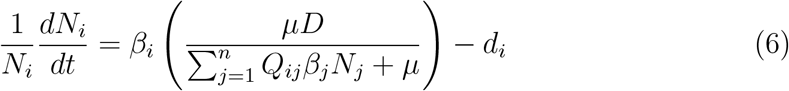

Finally, applying the first order Taylor expansion on 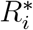 on equations (5) and (6) yields the classical Lotka-Volterra competition model (Figure 1D):

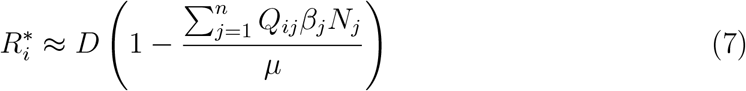

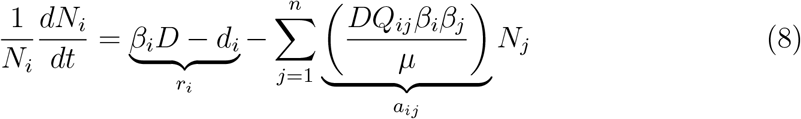

where the parameters of the original multistrain model can be rearranged into growth rates *r*_*i*_ and competition coefficients *a*_*ij*_.

An essential point is that this Lotka-Volterra competition model differs from the Lotka-Volterra predator-prey model. The former model typically assumes positive intrinsic growth rates *r*_*i*_ for all species, whereas the latter assumes a negative intrinsic growth rate *r*_*i*_ for the predator, which requires predation of prey species to persist (Volterra, 1927; Lotka, 1932). These differences permit cycles in both predator and prey populations, driven by the overcompensation of the resource (prey) population. Throughout the paper, we focus on the dynamics of the Lotka-Volterra competition model to characterize the differences in competitive structures between multistrain and ecological systems.

This conceptual framework for bridging an ecological competition model with an epidemiological strain competition model reveals two main discrepancies between the Lotka-Volterra model and the multistrain model. First, the Lotka-Volterra competition model implicitly assumes that the resource dynamics are always at equilibrium, whereas the multistrain model explicitly accounts for the susceptible host dynamics. Second, the Lotka-Volterra model underestimates equilibrium densities compared to the multistrain model, even though the nonlinear competition model that assumes equilibrium resource dynamics can seemingly predict the equilibrium accurately (Supplementary Figure S1). This underestimation of equilibrium densities indicate that there is yet another missing link between the multistrain model and the Lotka-Volterra model.

A comparison of resource dynamics across the three model formulations (i.e., multistrain model, nonlinear-competition model, and Lotka-Volterra model) reveals that the Lotka-Volterra model overestimates the resource density available to the invader during the initial invasion phase (Supplementary Figure S1). This overestimation causes an underestimation of resident density and an overestimation in the invasion growth rate of the invader (Supplementary Figure S1). Thus, for the Lotka–Volterra model to produce identical equilibrium resource densities to those in the multistrain model, the two competitors must coexist at lower equilibrium densities. This raises the question of whether and when the Lotka-Volterra model will be useful for predicting the competition outcome of a realistic multistrain competitive system, and if these discrepancies can be resolved via utilizing different parameterizations. Hereafter, we first consider the impact of relative speed between consumer and resource dynamics on competitive dynamics and return to the question of equilibrium dynamics later on.

### Relative speed between consumer and resource dynamics

As mentioned earlier, one of the main differences between the Lotka-Volterra model and the multistrain model is the relative speed between consumer and resource dynamics, where the Lotka-Volterra model implicitly assumes that the resource dynamics occur at a much faster time scale then consumer dynamics and therefore are always at quasi-equilibrium. To understand how the differences in time scale between consumer and resource dynamics influence competition, we explicitly model separate time scales for consumer and resource dynamics (*dC/dt* and *dR/dϵ*), where *ϵ* = *st* represents the relative speed of resource dynamics compared to consumer dynamics such that *s >* 1 implies that resource dynamics are faster than consumer dynamics. We then simulate stochastic competition between two species using randomly drawn parameters and assess the outcome of competition (Figure 2; Supplementary Text S3). We constrain the parameter space to always permit mutual invasion in order to focus on the role of resource overcompensation, rather than competitive exclusion, in preventing long-term coexistence.

**Figure 2:**
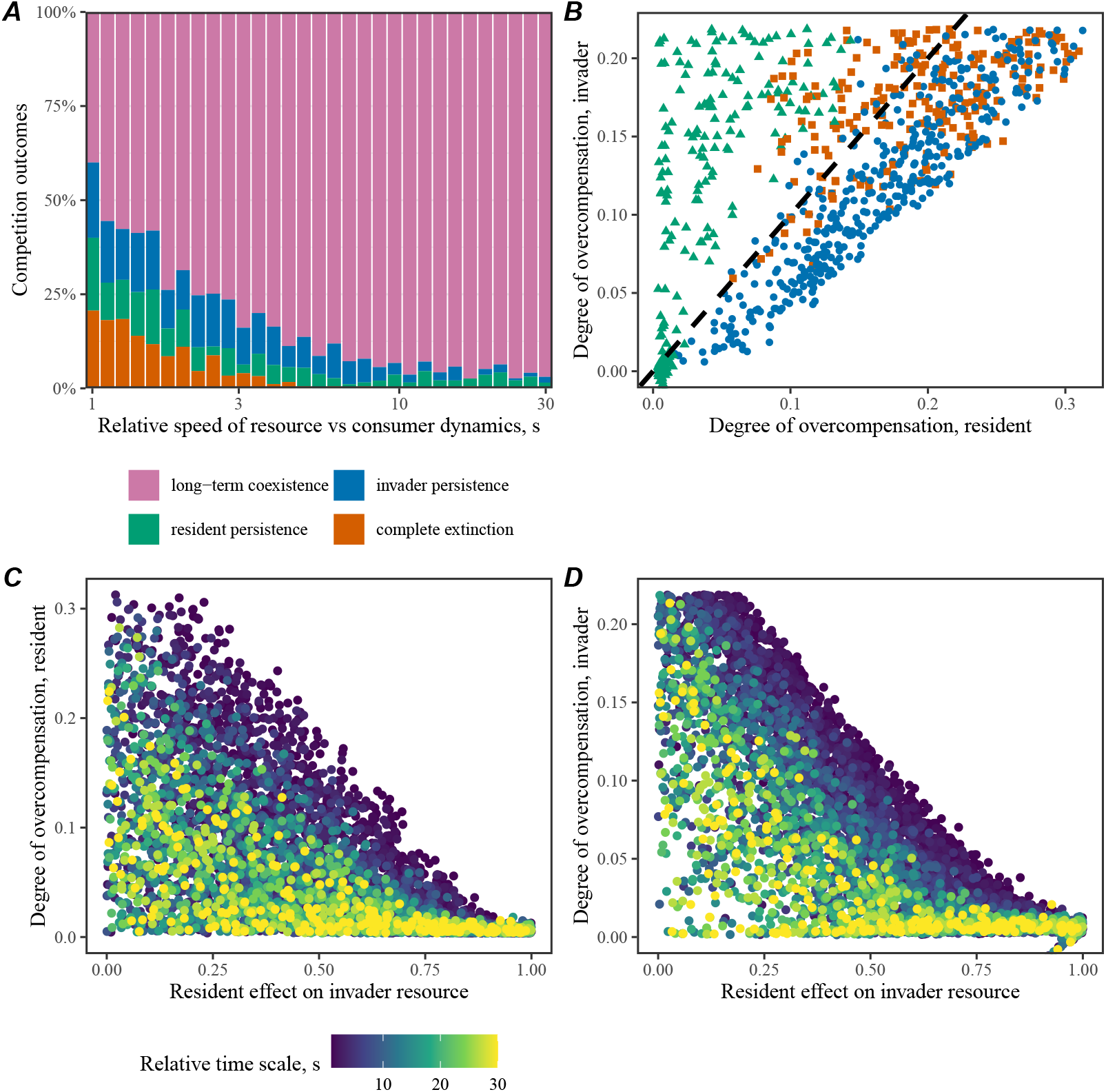
Relative speed between consumer and resource dynamics and the degree of overcompensation determines the outcome of stochastic competition. We simulated the multistrain model by separating the time scale of consumer and resource dynamics by a factor of *s* such that *s >* 1 implies faster resource dynamics across random parameters and evaluate the outcome of competition (long-term coexistence, invader persistence, resident persistence, or compete extinction). (A) Proportion of competition outcomes across different values of relative time scale of resource dynamics. (B) The relationship between the degree of overcompensation and resulting competition outcomes. The degree of overcompensation is calculated by taking the difference between the equilibrium resource proportion and minimum resource proportion. The dashed line represents the one-to-one line. Coexistence outcomes are not shown for visual clarity and are presented in Supplementary Figure S2 instead. (C–D) The relationship between the resident effect on invader resource and the degree of overcompensation.

We find that the relative speed between consumer and resource dynamics is a major determinant for long-term coexistence, where the probability of long-term coexistence given mutually invasible pairs of competitors becomes near 1 as we speed up the resource dynamics, *s* ≫ 1 (Figure 2A). In contrast, the majority of simulations did not result in long-term coexistence when consumer and resource dynamics were operating at similar speed, *s* ≈ 1. When two species fail to coexist, the outcome of their competition can be explained by the variation in the degree of resource overcompensation (Figure 2B). Whichever competitor experiences a greater resource overcompensation will fail to persist; in cases where both competitors experience large resource overcompensation, both competitors can become extinct. This degree of resource overcompensation is determined by the competition coefficient that a resident *r* imposes on the invader *i* (*Q*_*ir*_): weaker competition can permit a larger boom-bust cycle following an invasion, causing large resource overcompensation (Figure 2C,D). The relative speed between consumer and resource dynamics also affects the degree of overcompensation, where faster resource dynamics limits how much overcompensation can happen (Figure 2C,D). We note that resource overcompensation can still occur under rapid resource dynamics (Supplementary Figure S2); however, fast resource dynamics allow rapid replenishment of overcompensation and can therefore permit long-term coexistence even in the presence of resource overcompensation (Park et al., 2024).

Analyses of stochastic simulations highlight challenges in predicting species coexistence from a deterministic model under strong resource overcompensation. Indeed, other studies have also emphasized limitations in translating deterministic predictions about coexistence into competition outcomes for systems with finite populations, which can experience stochastic extinctions (Schreiber et al., 2023). This immediately raises the question of whether the simple Lotka-Volterra model can be still useful for understanding equilibrium dynamics of a multistrain system under limited resource overcompensation. For instance, a two competitor example in Figure 1 and Supplementary Figure S1 showed that the Lotka-Volterra model can still accurately predict the relative ordering of equilibrium densities despite the overall underestimation of equilibrium values. Thus, can this model still provide accurate predictions for the relative ordering of equilibrium densities for larger communities? Are there ways to address the apparent biases in predictions about equilibrium densities?

Thus, in the following sections, we systematically evaluate the ability of the Lotka-Volterra model to predict the equilibrium dynamics of more realistic multistrain systems and introduce a new mathematical framework for parameterizing the Lotka-Volterra model. In doing so, we introduce higher-order terms into the Lotka–Volterra framework, which have recently received increasing attention as key determinants of species coexistence (Letten and Stouffer, 2019; Gibbs et al., 2022; Kleinhesselink et al., 2022; Gibbs et al., 2024).

### Higher-order interactions determine the outcome of multistrain competition

To assess the accuracy of the Lotka-Volterra model for predicting the equilibrium dynamics, we need to be able to translate parameters of a general competition model into those of a Lotka-Volterra model. The first-order Taylor expansion that we showed earlier is one approach (Figure 1), but there are several limitations to this method. First, as we discussed earlier, this approach tends to underestimate equilibrium densities (Figure 1; Supplementary Figure S1)—these biases persist even for larger communities (compare Figure 3B and 3C). Moreover, the Lotka-Volterra model based on the first-order Taylor expansion around equilibrium fails to predict the ordering of equilibrium densities for larger systems (Figure 3F). Thus, while the derivation we presented in Figure 1 provides a useful conceptual linkage between ecological and epidemiological communities, its practical applications are limited. To address this gap, we develop a new mathematical theory for quantifying pairwise and higher-order interaction coefficients of Lotka Volterra model from any competition system (Figure 3A), and compare predictions from a multistrain parameterization for the consumer-resource model (Figure 3B) against those from Lotka-Volterra using different parameterizations (Figure 3C–H). This new paramterization is particularly useful because it can be applied to any general competition systems, beyond multistrain systems we present here; in contrast, the earlier parameterization relied on specific assumptions to derive Lotka-Volterra competition model from a multistrain system, and this expansion around equilibrium is infeasible for many models.

**Figure 3:**
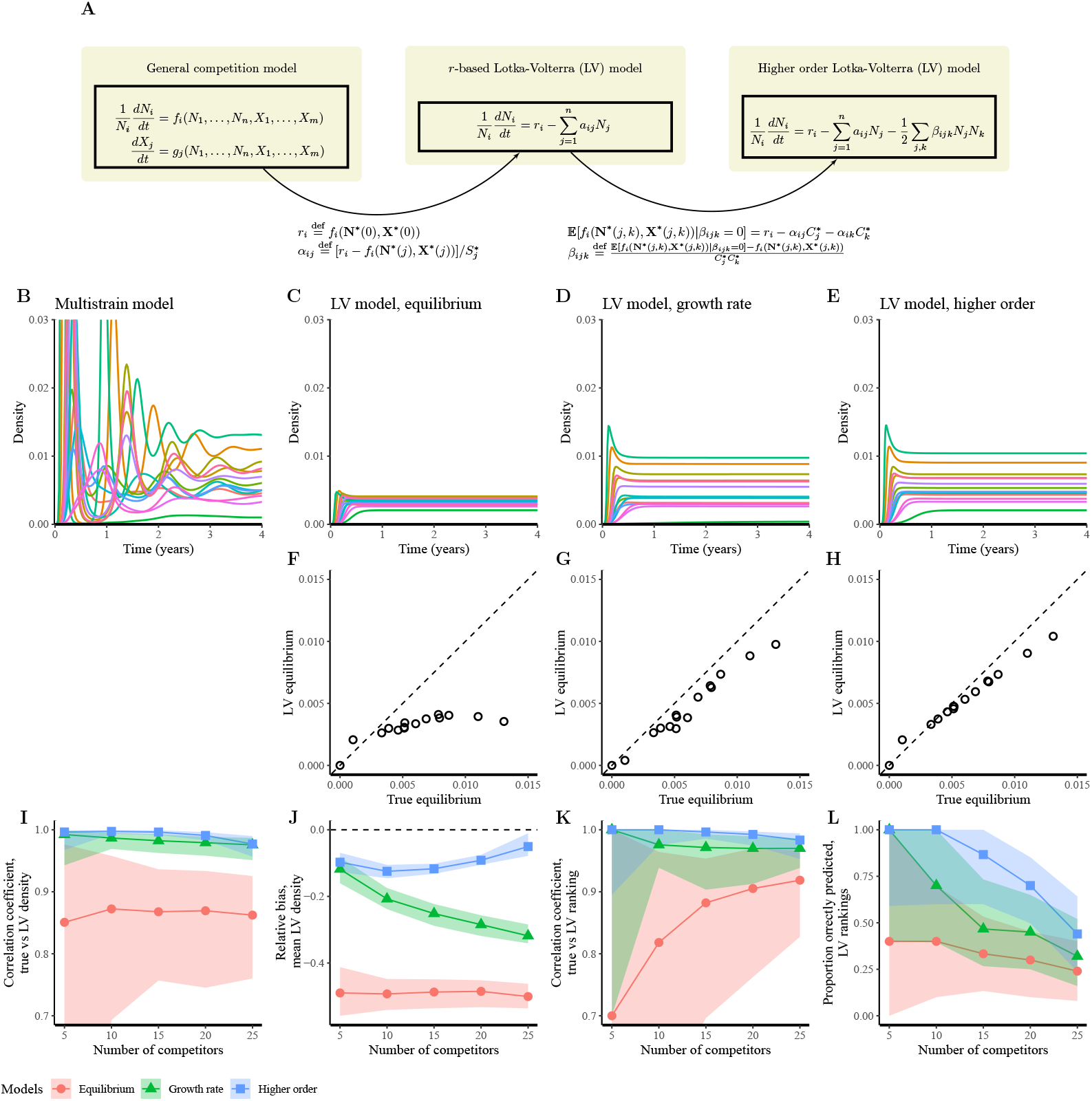
New framework for estimating pairwise and higher-order interaction coefficients allow accurate predictions for a consumer-resource model from simple Lotka-Volterra model. (A) Mathematical framework for estimating pairwise and higher-order interaction coefficients from any generic competition model. (B) Simulated trajectories of infection prevalence by each strain from a consumer-resource model based on the multistrain parameterization. (C–E) Corresponding Lotka-Volterra simulations based on (C) linearization at equilibrium, (D) parameterization from invasion growth rate, and (E) parameterization including higher-order interaction terms. (F–G) Comparisons between the true equilibrium based on a multistrain system and predictions based on Lotka Volterra models. (I) Correlation coefficients between the true and predicted equilibrium strain densities. (J) Mean relative bias between the true and predicted equilibrium strain densities. (K) Correlation coefficients between the true and predicted rank orders of equilibrium strain densities. (L) Proportion of strain rank orders of equilibrium strain densities correctly predicted by the Lotka–Volterra model.

To introduce a framework for quantifying pairwise and higher-order interaction coefficients, we begin with a generic expression capturing species competition:

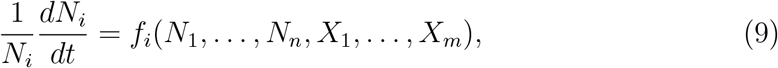

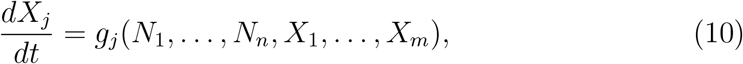

where *N*_*i*_ represents the density of species for *i* = 1, …, *n*; *X*_*i*_ represents the amount of modulator variables that shape the competition for *i* = 1, …, *m*; and *f*_*i*_ and *g*_*j*_ represent the per capita growth rate of species and modulator variables, respectively. Here, modulator variables *X*_*i*_ essentially captures all other variables that do not correspond to our main competitors *N*_*i*_, including resource variables but still affect the competition. When we consider a more realistic strain competition model later on, we focus on the density of primary infections caused by each strain *i* as our main density variables *N*_*i*_ and treat all other variables representing the dynamics of susceptible, recovered, and secondary/tertiary infections as modulator variables. Then, our framework seeks to quantify pairwise and higher-order interaction terms by (1) taking the difference between the invasion growth rate that we expect to observe in the absence of focal interaction and the realized growth rate that we observe when focal competitors are at equilibrium and (2) dividing this difference by the equilibrium density of the focal competitor. For example, the pairwise competition coefficient *α*_*ij*_ is obtained by comparing the intrinsic growth rate in the absence of any competitor *r*_*i*_ against the realized growth rate *f*_*i*_(**N**^∗^(*j*), **X**^∗^(*j*)) when competitor *j* is at equilibrium:

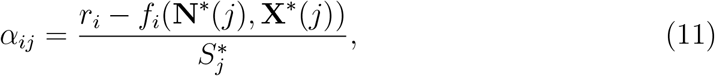

where the vector 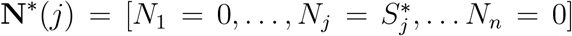 denotes the equilibrium when only species *j* is present; the vector **X**^∗^(*j*) denotes the corresponding equilibrium for modulator variables *X* for the condition **N**^∗^(*j*); and 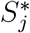 represents the single-species equilibrium. This implies that the intraspecific competition coefficient *α*_*ii*_ can be written as a ratio between intrinsic growth rate and the equilibrium density:

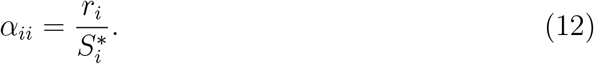

In other words, given any pairwise competition, this parameterization allows us to preserve the equilibrium density of the resident and the invasion growth rate of the invader. As we illustrate in Supplementary Text S4, this property also allows us to preserve the mutual invasibility of any two competitors and (2) derive a generic metric for niche and fitness differences that extend Chesson’s Modern Coexistence Theory (Chesson, 2000). As we further illustrate in Supplementary Text S5, the coexisting equilibrium densities for a pairwise competition system depends on the invasion growth rates, intrinsic growth rate, and single-species equilibrium density of the resident—since this new parameterization allows us to preserve all these quantities, we expect this parameterization to provide better predictions about equilibrium densities of larger communities than the earlier parameterization, which failed to preserve to resident densities and invasion growth rates.

Similarly, the higher-order interaction term *β*_*ijk*_ is obtained by comparing (1) the growth rate of species *i* that we would expect to observe in the absence of higher order interactions when species *j* and *k* are at equilibrium, 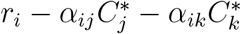, and the realized growth rate under the same condition, (2) *f*_*i*_(**N**^∗^(*j, k*), **X**^∗^(*j, k*)):

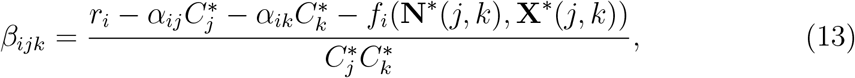

where 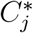 and 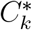 represent the equilibrium densities of species *j* and *k* when they are both at coexisting equilibrium. Parameterizing pairwise and higher-order interaction terms based on the differences in expected and realized invasion growth rates allows the Lotka-Volterra model to preserve invasion growth rates of more realistic competition model under the modeled interactions. Full details of mathematical derivations are presented in Supplementary Text S4.

As we mentioned earlier, the simple Lotka-Volterra model, derived from a firstorder Taylor approximation around the equilibrium (Figure 1C), fails to capture either the equilibrium density or the ordering of competitor densities (Figure 3C,F). Instead, when we calculate pairwise competition coefficients from invasion growth rates, the Lotka-Volterra model provides a much better prediction for the equilibrium density (Figure 3D,G). Inclusion of higher order interaction terms further improve predictions for equilibrium density and the ordering of competitor densities (Figure 3E,H).

We systematically evaluate the ability of a simple Lotka-Volterra model for predicting the deterministic outcome of multistrain competition by measuring (1) correlation coefficients between true and predicted equilibrium densities (Figure 3I), (2) relative biases in predicted equilibrium densities (Figure 3J), (3) correlation coefficients between the true and predicted rank orders of equilibrium densities (Figure 3K), and (4) proportions of strain rank orders of equilibrium strain densities that were correctly ordered (Figure 3L). These quantities are measured across randomly drawn parameters by increasing the number of competitors from 5 to 25. Detailed are provided in Supplementary Text S6.

Across all scenarios, the Lotka-Volterra model, parameterized from a first-order Taylor approximation around the equilibrium, performs poorly (Figure 3I–L). In contrast, the Lotka-Volterra model, based on the invasion growth rate analysis, excels in predicting the ordering equilibrium densities (Figure 3I,K). There are non-negligible biases in the predictions for equilibrium density values, but these biases are substantially reduced when we include higher order interaction terms (Figure 3J). However, predicting the exact ordering of the equilibrium densities for each competitor becomes more difficult as we increase the number of competitors (Figure 3L).

So how do higher order terms arise from a simple multistrain model even though the model only assumes pairwise interactions? In our simulations, more than 97% of estimated higher order terms are negative, implying that the realized invasion growth rate of a focal strain *i* when rare (when two competing strains *j* and *k* are at coexisting equilibrium) is higher than the invasion growth rate that we expect to observe in the absence of higher order terms (Supplementary Figure S3). In fact, this higher order term arises, not because of nonlinear interactions between strains *j* and *k*, but simply because of nonlinear susceptible depletion that strain *i* experiences (see Supplementary Text S7 for detailed mathematical explanation). Briefly, the Lotka-Volterra model implicitly assumes a linear effect of strains *j* and *k* on the equilibrium susceptible host density against strain *i*. However, under the explicit multistrain system, the equilibrium susceptible host density decreases nonlinearly as a function of density of competing strains *j* and *k*. Thus, a linear reduction in susceptible dynamics overestimates the amount of susceptible depletion that strain *i* will experience in the presence of strains *j* and *k*, thereby yielding negative higher order terms. Interestingly, we show analytically that the resulting higher order term for the growth rate of strain *i* is negative even in the absence of any interactions between strains *j* and *k*. Overall, these analyses highlight that nonlinear host-pathogen interactions can give rise to higher order interactions, which provide crucial information about understanding coexistence patterns in large pathogen strain communities.

### Case study: Rotavirus strain coexistence

Finally, we apply the resulting framework to rotavirus strain dynamics as a case study to test the utility of the Lotka Volterra approximation—in particular, we focus on how the cross immunity structure affects our ability to predict equilibrium dynamics of a realistic pathogen strain system from a simple ecological model. Rotavirus is one of the main causes of severe gastroenteritis and is responsible for a major childhood mortality (Estes et al., 1983; Glass et al., 2006). One of the main challenges in controlling rotavirus arise from serological variation in rotavirus strains, where vaccination may alter the strain distribution and provide opportunities for new strains to emerge (Pitzer et al., 2011). Previously, Pitzer et al. (2011) used a detailed mechanistic model, incorporating differences between homotypic vs heterotypic immunity as well as stage-structured infections, to understand rotavirus strain dynamics. Here, we ask whether a simple Lotka-Volterra model can accurately predict rotavirus strain distributions by applying our framework (Supplementary Text S8).

In order to apply our framework, we need to be able to quantify invasion growth rates, and therefore, we simplify the original model by excluding seasonal transmission (Supplementary Text S8). Even in the absence of seasonal transmission, the model predicts a cycle between competing strains (Figure 4A). In contrast to the original model, which assumed one dominant strain vs four sub-dominant strains with equal fitness, we allow all five strains to have different transmissibility to test the ability of a Lotka-Volterra model for accurately predicting the ordering of their fitness (Supplementary Text S8). Specifically, under our parameterization, only four strains can coexist simultaneously and the least transmissible strain becomes extinct (Figure 4B).

**Figure 4:**
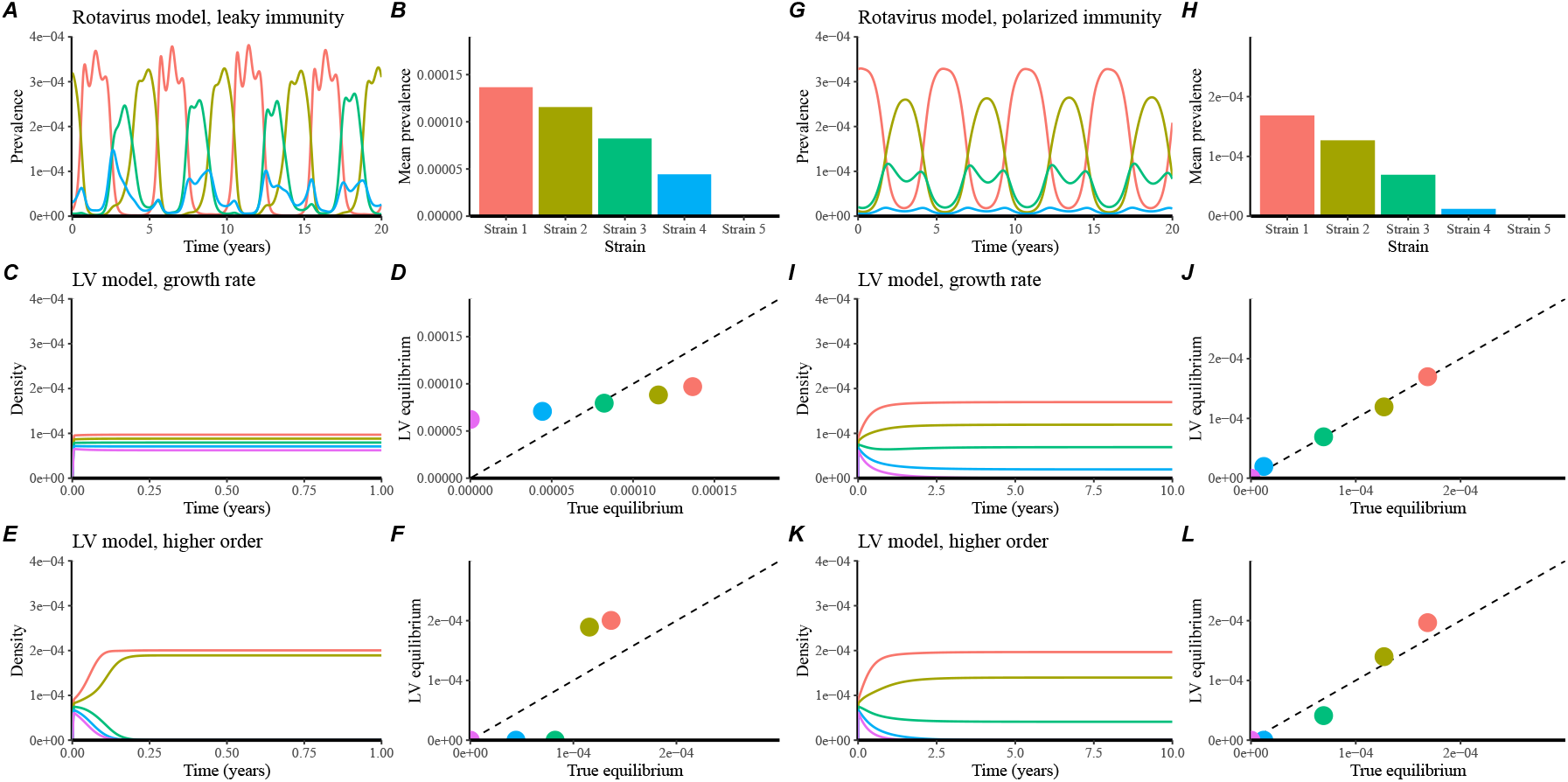
Comparisons between strain coexistence patterns from a mechanistic rotavirus strain competition model assuming leaky (A–F) and polarized (G–L) immunity vs Lotka-Volterra model. (A, G) Simulated trajectories of infection prevalence caused by each strain based on the mechanistic transmission model. (B, H) Mean prevalence of infection caused by each strain over 20 years. (C, I) Simulated trajectories of infection prevalence caused by each strain based on the Lotka Volterra model based on the invasion growth rate parameterization. (D, J) Comparisons between the true and predicted infection prevalence. Dashed lines represent the one-to-one line. (E, K) Simulated trajectories of infection prevalence caused by each strain based on the Lotka Volterra model accounting for higher order interaction terms. (F, L) Comparisons between the true and predicted infection prevalence. Dashed lines represent the one-to-one line. Across all panels, colors represent 5 different rotavirus strains.

Applying our framework to the rotavirus model, we find that the Lotka-Volterra model provides poor predictions for equilibrium prevalence for rotavirus strains (Figure 4C,D). The Lotka-Volterra model accurately predicts the ordering of strain prevalence but the exact equilibrium values are severely biased (Figure 4D). Even when we include higher order interactions, the Lotka-Volterra model fails to provide accurate predictions for equilibrium strain prevalence (Figure 4E,F).

So why does the Lotka-Volterra model perform so well in simulations (Figure 3) but poorly for this empirical system (Figure 4)? The answer lies in the differences in assumptions about underlying immunity. The multistrain model, derived from the consumer-resource model, assumes a polarized immunity, meaning that individuals are either fully susceptible or fully immune (Gog and Grenfell, 2002). In contrast, the rotavirus model by Pitzer et al. (2011) assumes leaky immunity, meaning that infection reduces susceptibility of all recovered individuals by the same, constant amount.

So we consider a polarized immunity formulation of the rotavirus model (Supplementary Text S8) and test whether the Lotka-Volterra model can predict the equilibrium prevalence for this model (Figure 4G–L). In this case, a simple Lotka-Volterra model with only pairwise competition provides a near perfect predictions for equilibrium densities (Figure 4I,J). In fact, including higher-order interactions make predictions slightly worse (Figure 4K,L). This case study illustrates that the underlying immunity structure, especially leaky vs polarized immunity, is a key determinant of strain competition that distinguishes host-pathogen systems from ecological communities.

## Discussion

There has been major theoretical progress in understanding species and strain coexistence across both community ecology (Allesina and Tang, 2012; Ellner et al., 2019; Gibbs et al., 2022; Yamamichi et al., 2022; Spaak and Schreiber, 2023; Miller et al., 2024) and infectious disease dynamics (Cobey and Lipsitch, 2012; Seabloom et al., 2015; Sieben et al., 2022; Park et al., 2024; Zilio et al., 2026). However, both fields have relied on disparate models for explaining coexistence patterns until recently (Sieben et al., 2022; Park et al., 2024) with limited insights into differences between ecological communities and pathogen strains. Here, we unify coexistence theory for ecological communities and pathogen strains by showing that a classic multistrain model and the Lotka-Volterra model can be derived from a consumer resource model. In doing so, we show that resource overcompensation, a main limiting factor that prevents strain coexistence, depends on the relative speed of resource vs consumer dynamics, explaining why resource overcompensation is so prevalent in host-pathogen systems, but not in ecological communities.

We also introduce a general theory for quantifying pairwise and higher order interactions for any competition model. A simulation study illustrates that equilibrium densities of a multistrain system can be accurately predicted from a simple Lotka-Volterra model by utilizing this theory. However, applying the resulting framework to a more realistic, rotavirus strain competition reveals that a Lotka-Volterra model can successfully predict the equilibrium densities of rotavirus strains only under polarized immunity, and not under leaky immunity. Overall, our study highlights the importance of characterizing the underlying susceptible resource dynamics as well as cross immunity structure for predicting strain coexistence patterns.

Recently, Sieben et al. (2022) and Park et al. (2024) both showed that modern coexistence theory from community ecology literature can be applied to host-pathogen systems, illustrating the feasibility of combining coexistence theory across two fields. The former study focused on a version of modern coexistence theory for decomposing invasion growth rates (Chesson, 1994, 2000; Ellner et al., 2019), whereas the latter study focused on a version for predicting mutual invasion based on niche and fitness differences (Godoy and Levine, 2014; Godoy et al., 2014; Kraft et al., 2015). In particular, Park et al. (2024) showed that the niche and fitness differences of two competing pathogens can be calculated based on their population-level susceptibility at equilibrium, which captures the strength of intraspecific and interspecific immune pressure. The framework presented here generalized the work of Park et al. (2024), by showing that intraspecific and interspecific competition coefficients of a Lotka Volterra model can be derived from any competition model—including but not limited to models of strain and pathogen competitions. The resulting framework allows the Lotka Volterra model to have identical invasion growth rates as the original model.

Our framework presented also allows us to quantify higher order interaction terms from any competition model. These higher order terms need to be interpreted with care because they do not necessarily reflect mechanistic higher order interactions. Instead, they represent phenomenological terms that capture deviation in invasion growth rates from the expected value when two other competitions are at equilibrium. Thus, as we showed in our rotavirus example, these higher order terms can also arise from nonlinear dynamics (Abrams, 1983). This interpretation deviates from the ecological definition of higher order terms, which represents a modification to the underlying pairwise interaction by a third competitor (Gibbs et al., 2022). Thus, an important avenue for future research is characterizing mechanisms that give rise to such higher order terms (Letten and Stouffer, 2019; Kleinhesselink et al., 2022; Gibbs et al., 2024).

There are several limitations to our analysis. First, the framework we present here for quantifying pairwise and higher order interaction terms has only been tested against two systems: a consumer-resource model based on a multistrain parameterization and a more realistic, rotavirus model. While these examples may appear to lean heavily towards host-pathogen interactions, the status-based multistrain model that we used is a special case of a consumer resource model and shares a similar structure and parameterization that Letten and Stouffer (2019) used as a mechanistic basis for ecological higher order interactions. Thus, we feel that the examples provided here represent both ecological and epidemiological communities; nonetheless, applications to other, realistic systems are needed. Moreover, the estimation of pairwise and higher order interaction terms depend on equilibrium densities, which are rarely observed in real systems. While direct application of the resulting framework to empirical systems may be limited, as we showed in the rotavirus example, such empirical applications first require an independent parameterization of a mechanistic model, which can then provide equilibrium conditions through simulations. Finally, our framework focuses on invasion at equilibrium, but many empirical systems exhibit complex cycles (Abrams and Holt, 2002; Pitzer et al., 2011; Benincà et al., 2015; Yamamichi et al., 2022; Clark et al., 2026). For the analysis of rotavirus strain dynamics, we were able to still obtain invasion growth rates by removing seasonal forcing and speeding up the resource dynamics, but this approach is necessarily approximate. Future studies should consider expanding this framework based on long-term averages of invasion growth rates across different conditions (Ellner et al., 2019). Despite these limitations, our study presents a theoretical foundation for linking mechanistic models of species competition with phenomenological models, with potential applications beyond host-pathogen systems.

In conclusion, we present a unifying framework for explaining the differences between species and strain competition through the lens of consumer-resource dynamics and further illustrate the feasibility of predicting the outcome of deterministic strain competition from ecological models. In doing so, our results underscore the importance of a detailed understanding of resource dynamics for predicting species and strain persistence and therefore coexistence. More broadly, explicit synthesis of ecological and epidemiological theory is needed to uncover shared links between two processes and generate new mechanistic insights (Anderson and May, 1991; Grenfell, 2001; Grenfell et al., 2002; Cobey and Lipsitch, 2012; Fraser et al., 2014; Brett et al., 2017; Sieben et al., 2022; Park et al., 2024, 2026).

## Acknowledgements

We thank Chuliang Song for the helpful discussion. S.W.P. was supported by the New Faculty Startup Fund from Seoul National University and by the National Research Foundation of Korea (NRF) grant funded by the Korea government (MSIT) (RS-2026-25474574). J.M.L. was supported by the National Science Foundation (DEB-2022213). B.T.G. was suppored by the Princeton Catalysis Initiative and Princeton Precision Health. B.T.G. have been funded in whole or in part with Federal funds from the National Cancer Institute, NIH, under Prime Contract No. 75N91019D00024, Task Order No. 75N91023F00016. The content of this publication does not necessarily reflect the views or policies of the Department of Health and Human Services, nor does mention of trade names, commercial products, or organizations imply endorsement by the U.S. Government.

## Competing interests

The authors declare no competing interest.

## Data availability

All data and code are stored in a publicly available GitHub repository (https://github.com/parklab-snu/consumer-resource-coexistence).

## Supplementary Figure

**Figure S1:**
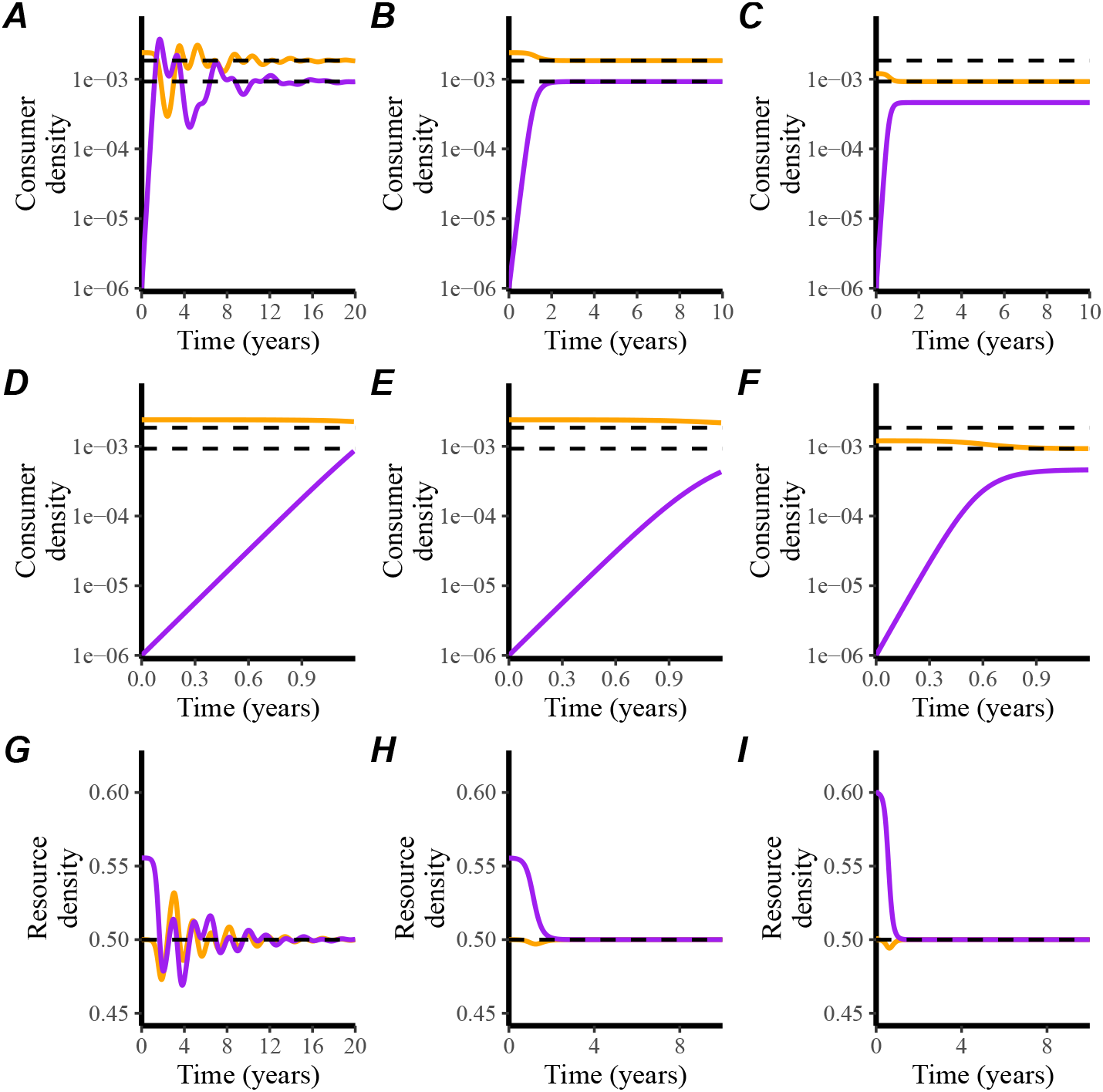
A side-by-side comparison of consumer dynamics between across different models. (A) Simulated trajectories of consumer dynamics from a multi-strain model. (B) Simulated trajectories of consumer dynamics from a non-linear competition model. (C) Simulated trajectories of consumer dynamics from the Lotka-Volterra model. (D–F) Zoomed-in views of the corresponding consumer dynamics plots in panels A–C. (G) Simulated trajectories of resource dynamics from a multi-strain model. (H) Simulated trajectories of resource dynamics from a non-linear competition model. (I) Simulated trajectories of resource dynamics from the Lotka-Volterra model. Dashed lines represent the equilibrium density values of the multi-strain model. These simulations are identical to the ones presented in Figure 1 in the main text.

**Figure S2:**
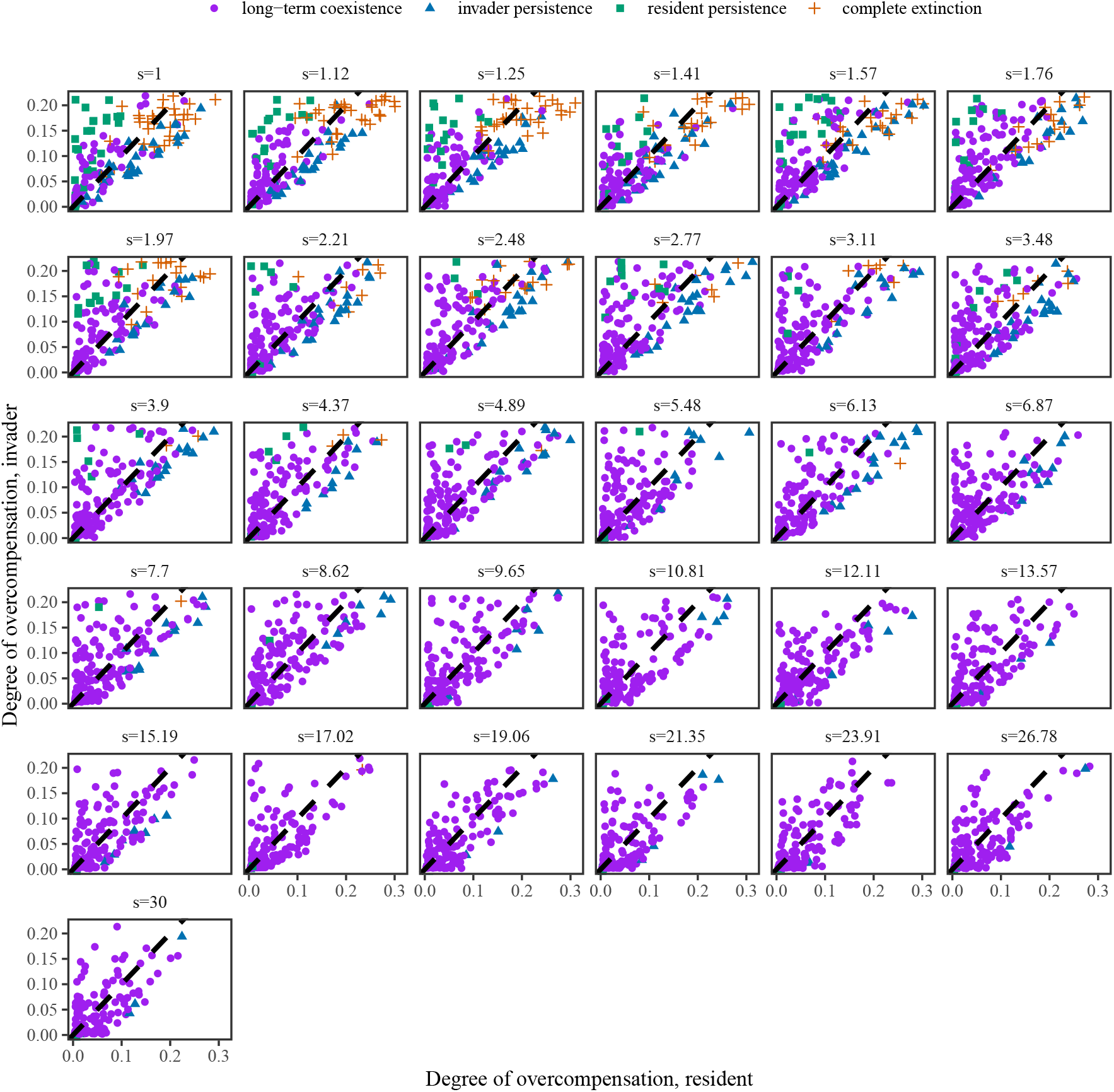
The relationship between the degree of overcompensation and resulting competition outcomes across different assumptions about the relative speed of resource dynamics. The relationship between the degree of overcompensation and resulting competition outcomes. The degree of overcompensation is calculated by taking the difference between the equilibrium resource proportion and minimum resource proportion. The dashed line represents the one-to-one line. Each panel corresponds to a value of the relative speed of resource dynamics *s*.

**Figure S3:**
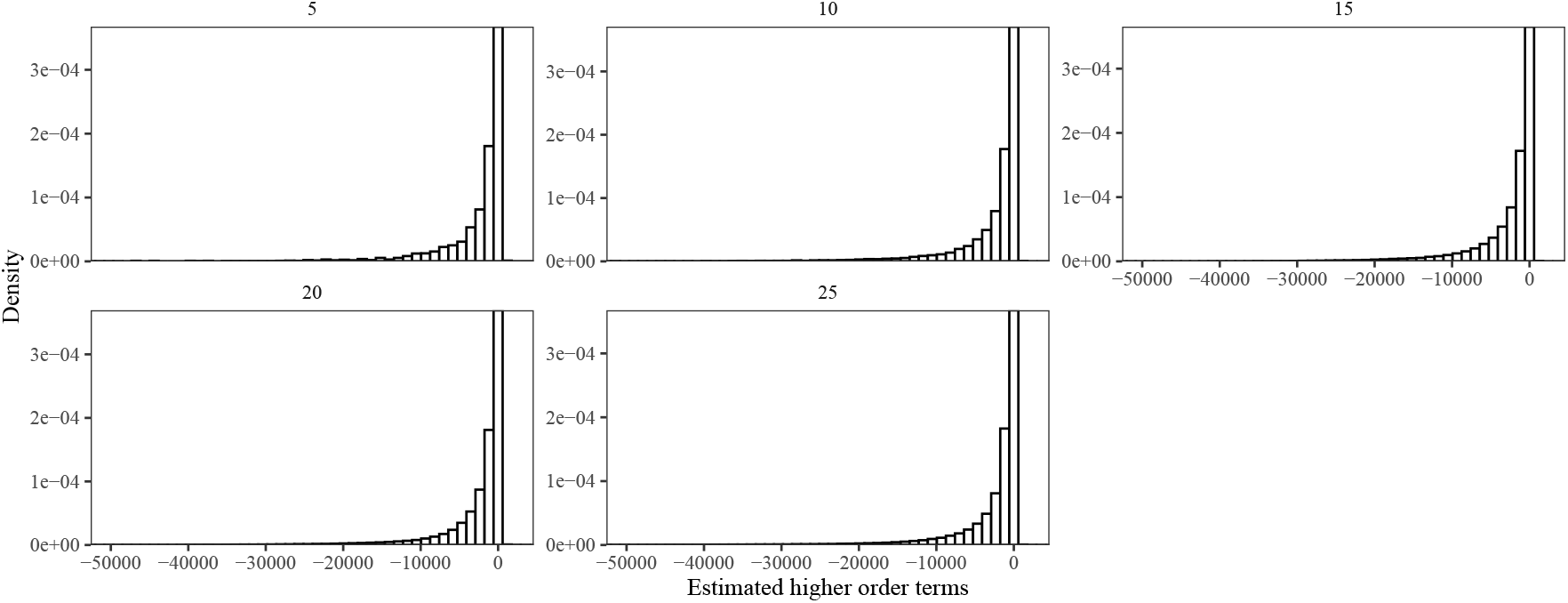
Estimated higher order terms from multistrain simulations. The histograms represent the distribution of estimated higher order terms from deterministic multistrain simulations. Each panel corresponds to the number of competitors in a multistrain system.

## Supplementary Text

### S1 Derivations of multistrain Lotka-Volterra model from the consumer-resource model

As discussed in the main text, we begin with a general consumer-resource model for *n* consumers *N*_*i*_ competing for *n* resources *R*_*j*_:

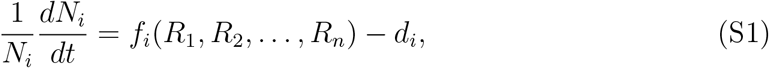

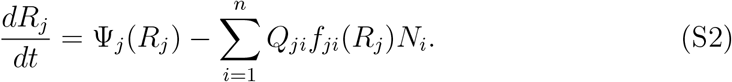

Under the set of assumptions laid out in the main text, the consumer-resource model simplifies to (Figure 1B):

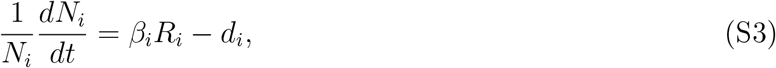

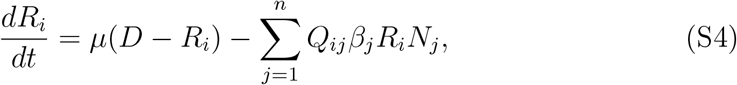

which corresponds to a status-based model for multistrain dynamics (Gog and Grenfell, 2002). Specifically, if we use the susceptible *S*_*i*_ to denote the density of susceptible hosts (resource) and *I*_*i*_ to denote the density of infected hosts (consumer), we obtain:

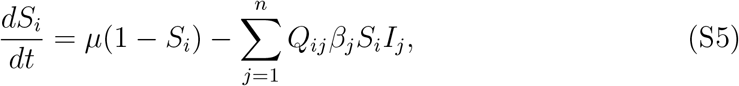

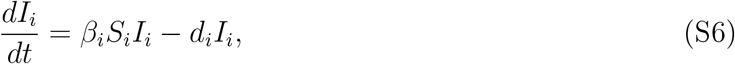

where *D* = 1 represents the assumed population size, and 1*/d*_*i*_ represents the mean infectious period (Gog and Grenfell, 2002).

Assuming that the resource dynamics is always at equilibrium, the competition model simplifies to::

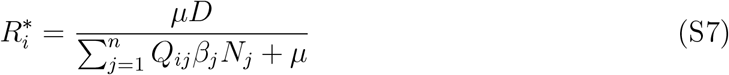

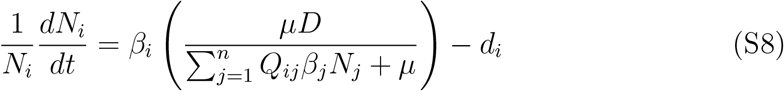

Finally, to further simplify this, we begin by writing

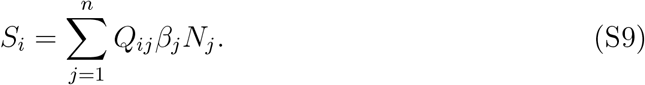

Then, we have

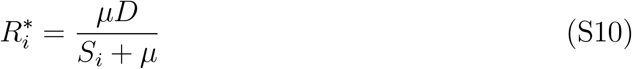

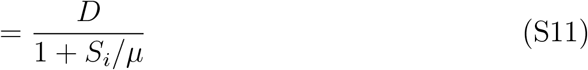

Assuming *S*_*i*_ ≪ *µ*, we can apply the first order Taylor approximation on 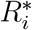:

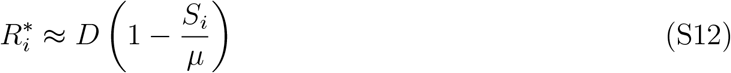

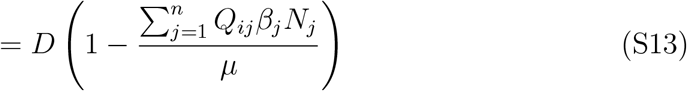

Then, the competition model simplifies to the classical Lotka-Volterra model:

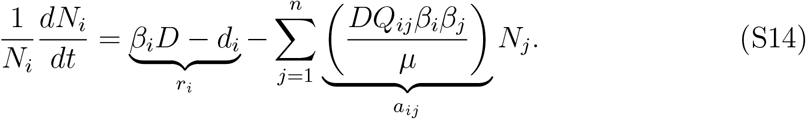

### S2 Waning immunity in multistrain model

The original status-based model by Gog and Grenfell (2002) assumes a life-long immunity:

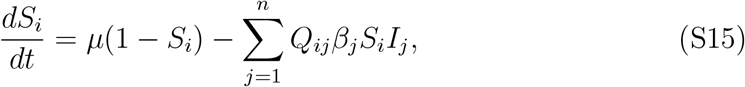

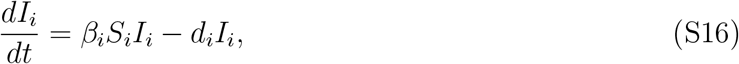

where the susceptible recruitment is driven by birth at rate *µ*. Instead, it is possible to include immune waning by imposing an SIRS structure:

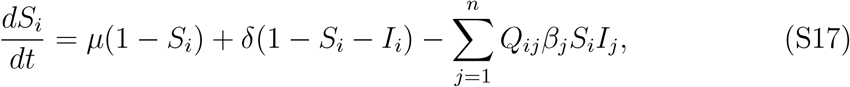

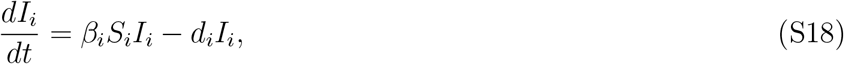

where *δ* represents the immune waning rate (Yang et al., 2020). In practice, the density of infected individuals are far smaller than the density of susceptible individuals (*I*_*i*_ ≪ *S*_*i*_). Therefore, assuming 1 − *S*_*i*_ − *I*_*i*_ ≈ 1 − *S*_*i*_, we can write:

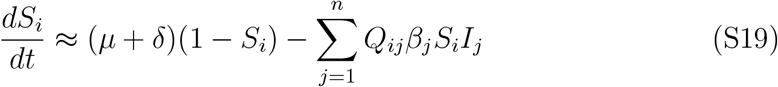

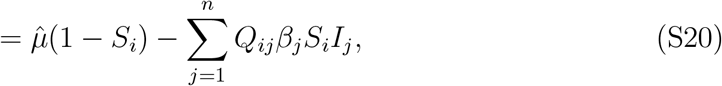

where 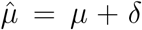 represents the effective rate of susceptible recruitment/immune waning. In other words, increasing the birth rate in the original status-based model can effectively capture the dynamics of immune waning and its implications for resource overcompensation and long-term coexistence.

### S3 Stochastic simulation of two competitors

To understand how the relative speed between consumer and resource dynamics affect the outcome of competition, we first begin by separating the time scale of consumer and resource dynamics:

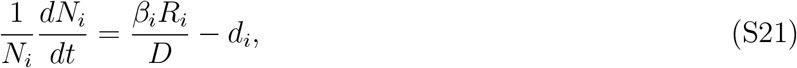

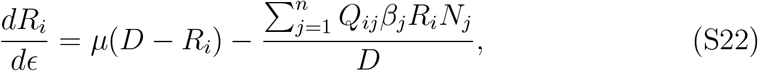

where *ϵ* represents the resource time scale. Writing *ϵ* = *st*, we have:

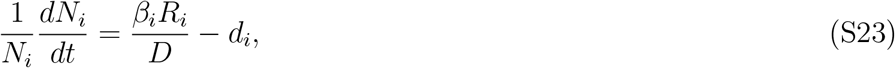

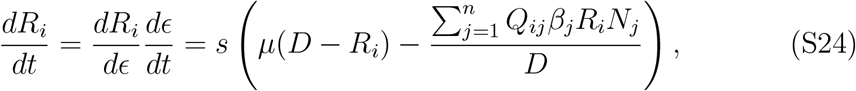

where *s* represents the relative time scale of resource dynamics compared to consumers with *s >* 1 representing a faster time scale of resource dynamics. We implement a stochastic version of this model to allow resource overcompensation to drive population extinction, using a binomial Euler scheme (He et al., 2010):

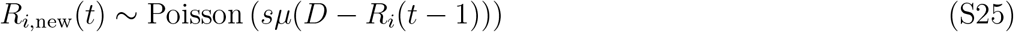

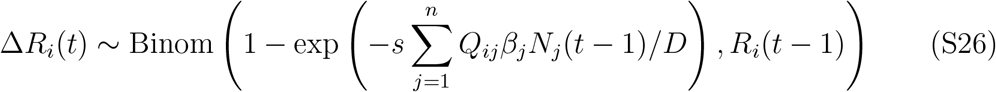

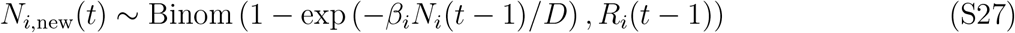

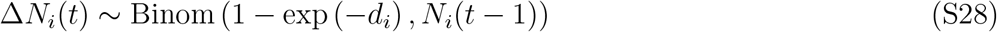

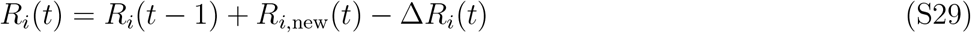

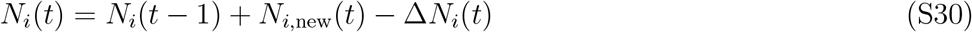

Then, we simulate the stochastic model across a plausible range of parameters for respiratory pathogens and evaluate the outcome of competition as well as the degree of overcompensation. Specifically, we vary *s* between 1 and 30 using 31 equi-valued steps on a log scale. For each value of *s*, we perform 200 competition simulations where the duration of immunity 1*/µ* is drawn from a gamma distribution with a mean of 5 years and a shape parameter of 2 (95% quantile: 0.6—13.9 years) and the basic reproduction number minus 1 (ℛ_0_ −1) is drawn from a gamma distribution with a mean of 1 and a shape parameter of 5 (95% quantile: 0.3—2.0):

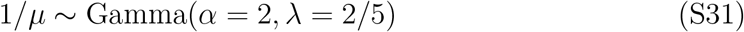

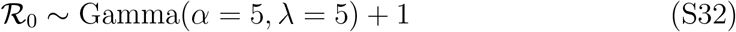

For simplicity, we assume that both competitors have the same ℛ_0_. Then, assuming a mean infectious period of 7 days (1*/d*_*i*_ = 7 days), we have: *β*_*i*_ = ℛ_0_*d*_*i*_ for both *i* = 1, 2. Finally, we assume *Q*_11_ = *Q*_22_ = 1 and draw *Q*_12_ and *Q*_21_ from independent uniform distributions between 0 and 1. Since two competing pathogens have identical ℛ_0_ and imperfect cross immunity, mutual invasion is always possible (Park et al., 2024). For each parameter set, we first simulate the model for 20 years using only the first competitor, where the number of susceptible and infected host are initialized at the continuous-time equilibrium with a population size *D* of 10^7^; this initial step allows the system to reach equilibrium and account for discrepancy between continuous- and discrete-time models. Then, we introduce 100 infected individuals for the second competitor and simulate the model for 20 additional years and assess which competitors have non-zero density at the end of the simulation. We further characterize the degree of overcompensation of each competitor by measuring the difference between the equilibrium resource density and the minimum resource density over the final 20 years.

### S4 Mathematical framework for quantifying pairwise and higherorder interactions

Our goal is to derive a generalized framework for quantifying pairwise and higherorder interactions for a general model of species competition:

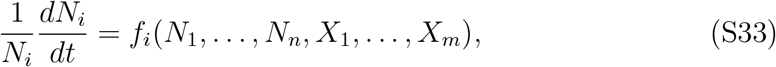

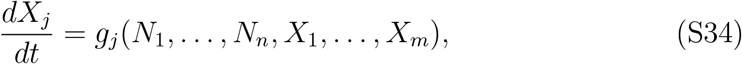

where *N*_*i*_ represents the density of species for *i* = 1, …, *n*; *X*_*i*_ represents the density of modulator variables that shape the competition for *i* = 1, …, *m*; and *f*_*i*_ and *g*_*j*_ represent the per capita growth rate of species and modulator variables, respectively. To derive the framework, we begin by considering a Lotka-Volterra model that includes higher-order interaction terms:

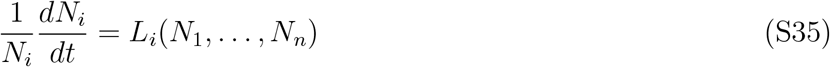

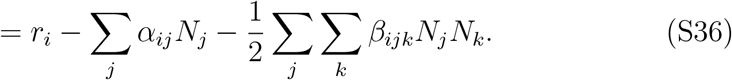

Here, we divide the higher order terms by 2 to avoid double counting of *β*_*ijk*_ and *β*_*ikj*_. This Lotka-Volterra model will provide a basis for calculating pairwise and higherorder interactions based on the differences in invasion growth rates across different conditions, divided by the density of corresponding competing species density at equilibrium.

First, the intrinsic growth rate *r*_*i*_ of species *i* represents the growth rate when rare in the absence of any competitors. From the Lotka Volterra model, we see that *r*_*i*_ can be expressed as *r*_*i*_ = *L*_*i*_(**N**^∗^(0)), where the vector **N**^∗^(0) = [*N*_1_ = 0, …, *N*_*n*_ = 0] denotes the equilibrium in the absence of any competitors. Analogously, we can also compute the growth rate when rare in the absence of any competitors for our model in Eq. (S33)–Eq. (S34):

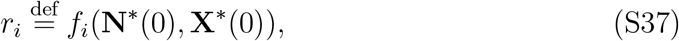

where the vector **X**^∗^(0) denotes the corresponding equilibrium for modulator variables *X* for the condition **N**^∗^(0).

Second, the intraspecific competition coefficient *α*_*ij*_ represents the per capita effect of species *j* in limiting the growth of species *i*. Rearranging the Lotka-Volterra equation, we can see that *α*_*ij*_ can be calculated by taking the difference between the intrinsic growth rate *r*_*i*_ and the invasion growth rate *f*_*i*_(**N**^∗^(*j*)) of species *i* when species *j* is at single-species equilibrium 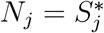, and further dividing this difference by the equilibrium density 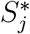:

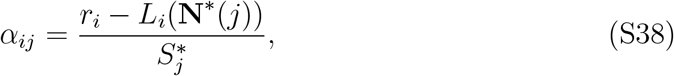

where the vector 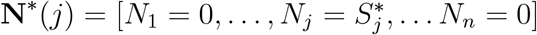 denotes the equilibrium when only species *j* is present. We use 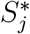 to describe the single species equilibrium so that it can be distinguished from equilibrium conditions when many coexisting species are present. Based on this observation, we define the intraspecific coefficient for the general model (Eq. (S33)–Eq. (S34)) to be:

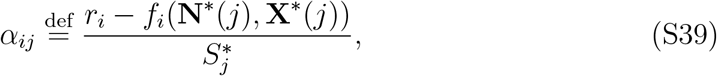

where the vector **X**^∗^(*j*) denotes the corresponding equilibrium for modulator variables *X* for the condition **N**^∗^(*j*).

Now, we can also define the intraspecific competition coefficient *α*_*ii*_ in a similar way:

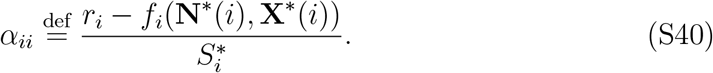

Since the invasion growth rate of species *i* is equal to zero when species *i* is at equilibrium (*f*_*i*_(**N**^∗^(*i*), **X**^∗^(*i*)) = 0), this allows us to simplify the expression for the intraspecific competition coefficient:

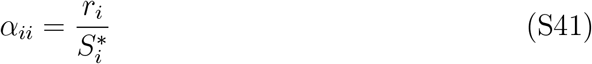

Finally, to quantify higher-order interactions, we begin by observing that the expected invasion growth rate of species *i* for Lotka-Volterra model when species *j* and *k* are at coexisting equilibrium (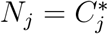 and 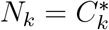) corresponds to:

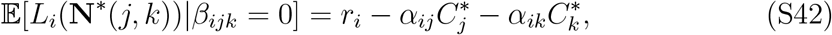

where the vector 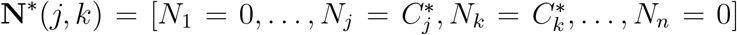 denotes the equilibrium when species *j* and *k* are at coexisting equilibrium. In contrast, in the presence of higher-order interactions, the invasion growth rate of species *i* corresponds to:

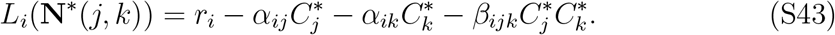

This suggests that higher-order interactions can be computed by taking the difference between the expected and observed invasion growth rates, and dividing this difference by the equilibrium densities of competing species 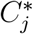 and 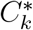:

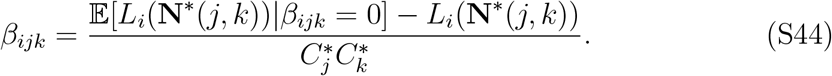

Therefore, for the general model of species competition (Eq. (S33)–Eq. (S34)), we can define the higher-order interaction term as follows:

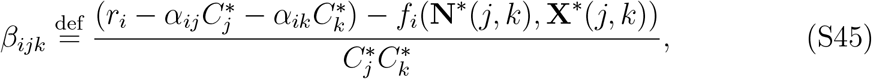

where the vector **X**^∗^(*j, k*) denotes the corresponding equilibrium for modulator variables *X* for the condition **N**^∗^(*j, k*). Here, the invasion growth rate *r*_*i*_ (Eq. (S37)) and interspecific competition coefficients *α*_*ij*_ and *α*_*ik*_ (Eq. (S39)) follow previous definitions.

Our framework now provides a natural way of defining niche and fitness differences for the general model of species competition (Eq. (S33)–Eq. (S34)). Specifically, based on intraspecific (Eq. (S40)) and interspecific competition (Eq. (S39)) coefficients that we defined above for a general model, we can define niche difference 𝒩 = 1 − *ρ* based on the ratio of geometric mean of the interspecific competition coefficient 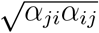 versus intraspecific competition coefficient 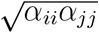

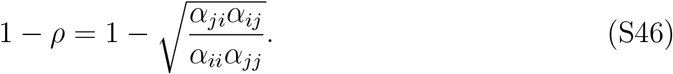

Likewise, we can define the fitness difference *f*_*j*_*/f*_*i*_ as a product of (1) the ratio between single species equilibrium (*S*_*j*_*/S*_*i*_) and (2) the ratio between geometric mean of per capita interspecific and intraspecific competition imposed by species *j, α*_*ij*_*α*_*jj*_, versus the corresponding value by species *i, α*_*ij*_*α*_*ii*_:

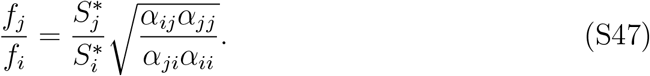

Then, it follows that the mutual invasion conditions for the general model (Eq. (S33)– Eq. (S34)) can be expressed as the following inequality:

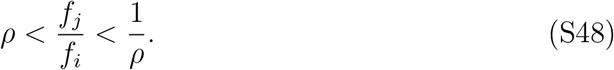

### S5 Relationship between invasion growth rates and coexisting equilibrium

Here, we explore the relationship between invasion growth rates and coexisting equilibrium for a simple Lotka-Volterra competition model. First, a pairwise Lotka-Volterra competition model can be written as:

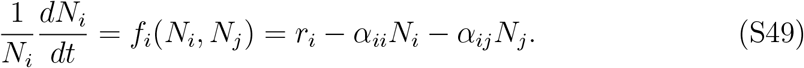

For this system, invasion growth rate 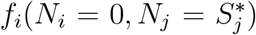 of species *i* when *j* is at equilibrium 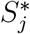 can be written as:

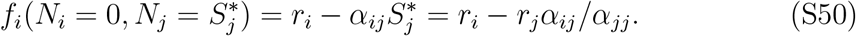

Likewise, we can also write down the invasion growth rate of species *j* when *i* is at equilibrium:

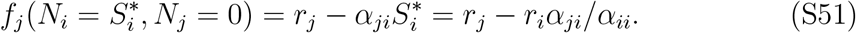

The coexisting equilibrium densities, 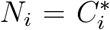 and 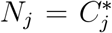, can be derived by solving the following set of equations:

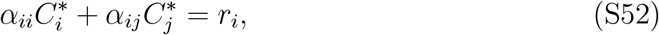

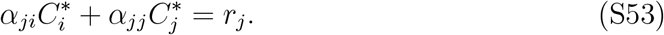

Solving this equation yields:

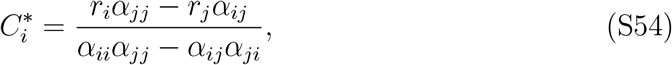

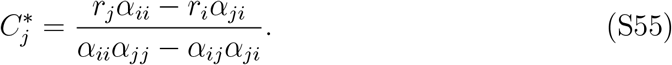

Then, this can be re-written as:

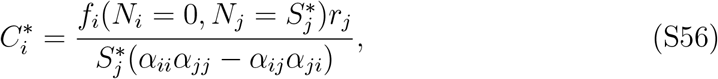

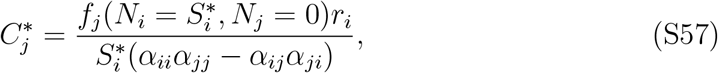

which shows that the coexisting equilibrium density of a focal species *i* is proportional to its own invasion growth rates 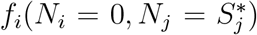 and the intrinsic growth rate of the competitor *r*_*j*_, and inversely proportional to the single-species equilibrium density of the competitor 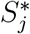. Thus, biases in any of these three quantities will lead to biases in predictions about coexisting equilibrium density.

### S6 Deterministic simulation of many competitors

We use deterministic simulations to assess the ability of Lotka-Volterra models to predict the equilibrium dynamics of a multistrain model. As described earlier, the model can be written as

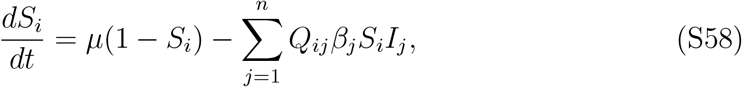

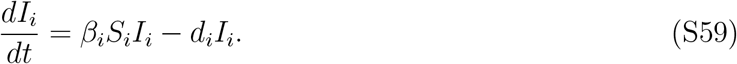

To test our framework, we vary the number of competing strains from 5 to 25, perform 100 simulations for each strain dimension using randomly sampled parameters, and evaluate the ability to accurately predict the equilibrium density. For each simulation, we sample the mean duration of immunity *D*_*I*_ and the basic reproduction number ℛ_0,*i*_ for each strain as follows:

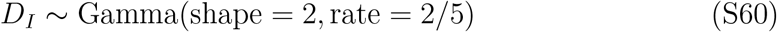

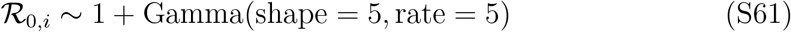

This allows the mean duration of immunity to have an average of 5 years (95% quantile: 0.6 years–13.9 years) and the basic reproduction number to have an average of 2 (95% quantile: 1.3–3.0)—these parameters are broadly consistent with many acute, respiratory pathogens. Assuming the average infectious period of 1 week (1*/d*_*i*_ = 1 week for all *i*), we have:

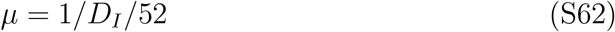

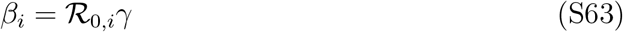

where *µ* represents the effective susceptible recruitment rate and *β*_*i*_ represents the transmission rate of strain *i*. The equilibrium values for each strain, in the absence of any competitors, can be calculated analytically:

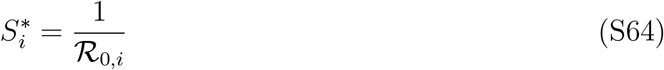

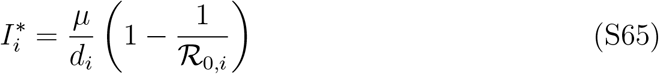

Then, non-diagonal entries of *Q*_*ij*_ are independently drawn from a uniform distribution between 0 and 0.1, with diagonal entries assumed to equal 1 (*Q*_*ii*_ = 1). Using these parameters, we simulate the model for 100 years, assuming a fully susceptible population and an initial infected fraction of 10^*−*6^ for each strain, so that the system can reach the equilibrium. Using the same parameters, we also simulate the Lotka Volterra model, linearized at equilibrium, to obtain the equilibrium density:

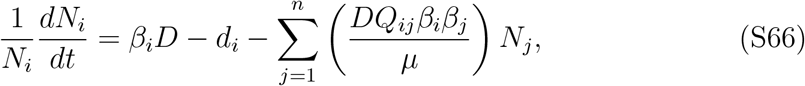

where *N*_*i*_ represents the density of infected population *I*_*i*_ with each strain.

We also consider alternative parameterizations for the Lotka Volterra model. First, we derive intraspecific and interspecific competition coefficients based on the analytical invasion growth rates. Specifically, the intrinsic growth rate of strain *i* for the multistrain model corresponds to

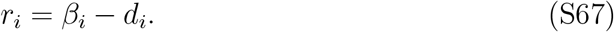

Then, the intraspecific competition coefficient is equal to:

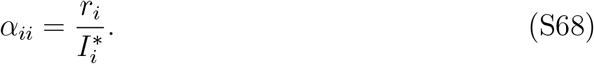

Note that when strain *j* is at equilibrium, the equilibrium susceptible fraction for strain *i* corresponds to:

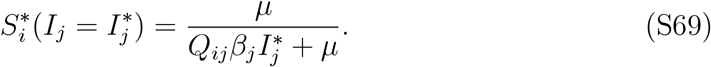

Thus, the interspecific competition coefficient is equal to:

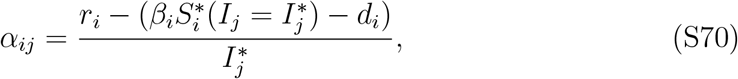

where 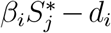 represents the growth rate of strain *i* when strain *j* is at equilibrium. Second, we compute the higher-order interaction terms based on analytical invasion growth rates. Specifically, strains *j* and *k* can coexist when

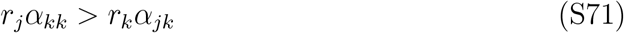

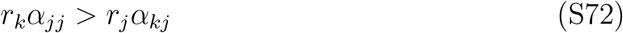

Under these conditions, the equilibrium susceptible fraction for these two strains still correspond to the following:

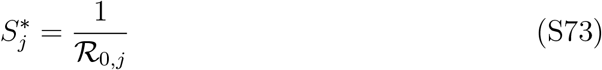

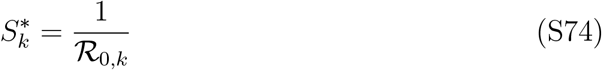

Thus, their coexisting equilibrium prevalence (*I*_*j*_ = *C*_*j*_ and *I*_*k*_ = *C*_*k*_) can be calculated by solving the following set of linear equations:

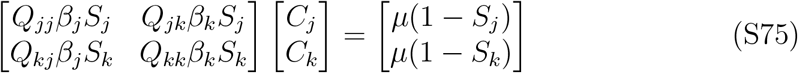

Then, the equilibrium fraction susceptible for strain *i* corresponds to:

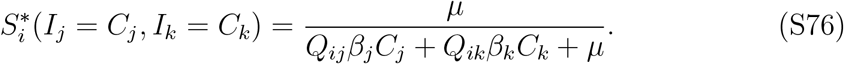

The expected growth rate of strain *i* equals:

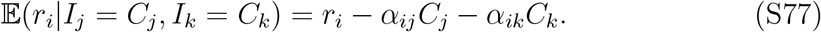

In contrast, the realized growth rate of strain *i* equals:

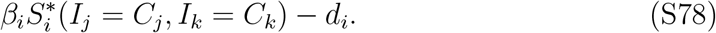

Therefore, the higher order interaction term corresponds to:

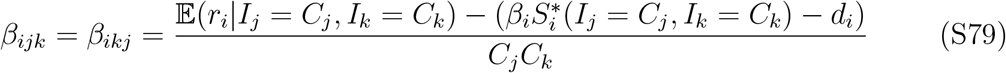

For each multi-strain simulation, we calculate the equilibrium density from three different Lotka-Volterra models and compare the accuracy of each model parameterization in predicting the true equilibrium densities.

### S7 Higher order terms in multistrain system

Estimating Lotka-Volterra coefficients from the multistrain system yields negative higher order terms across the majority of our simulations. To understand this phenomenon, we first consider the definition of higher order terms:

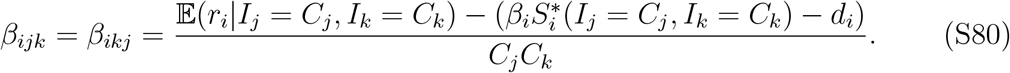

Here, higher-order terms *β*_*ijk*_ and *β*_*ikj*_ are defined based on the difference between the expected invasion growth rate, 𝔼(*r*_*i*_|*I*_*j*_ = *C*_*j*_, *I*_*k*_ = *C*_*k*_), and the realized invasion growth rate, 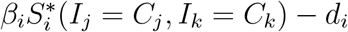.

The expected invasion growth rate under the pairwise Lotka-Volterra formulation corresponds to:

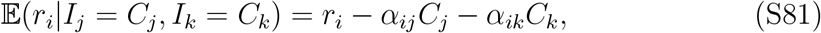

which assumes linear effect of competitors *j* and *k* on the invasion growth rate of competitor *i*. In contrast, the realized invasion growth rate corresponds to:

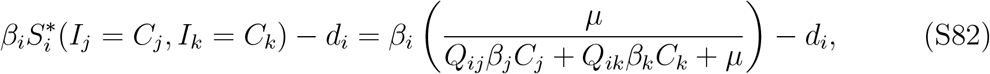

which assumes a nonlinear effect of competitors *j* and *k* on the invasion growth rate of competitor *i*.

To make the comparison more explicit, we begin by writing *δ*_*ij*_ = *Q*_*ij*_*β*_*j*_ and *δ*_*ik*_ = *Q*_*ik*_*β*_*k*_. Then, the pairwise interaction term *α*_*ij*_ can be written as:

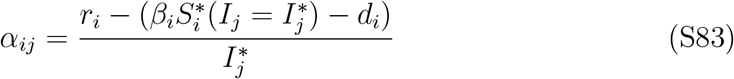

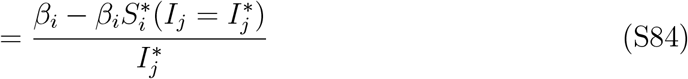

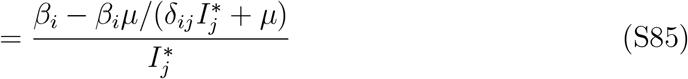

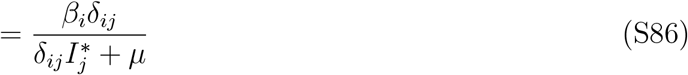

Analogously, we have:

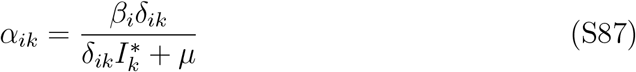

Then, the expected growth rate can be written as:

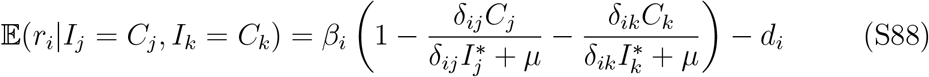

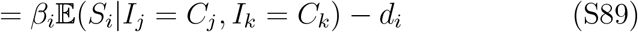

where 𝔼(*S*_*i*_ | *I*_*j*_ = *C*_*j*_, *I*_*k*_ = *C*_*k*_) represents the expected level of susceptible host density under linear effects of competitors *j* and *k*.

In contrast, the actual level of susceptible host density for the multistrain model is nonlinear:

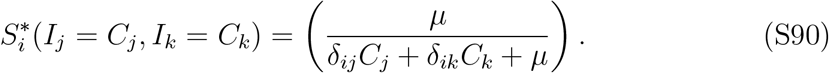

In particular, the differences in expected and realized invasion growth rate can be understood in terms of the differences in expected and realized susceptible host density:

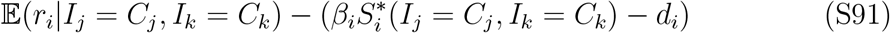

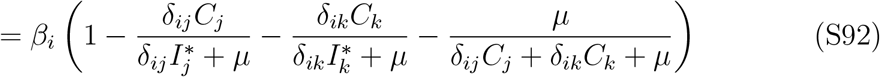

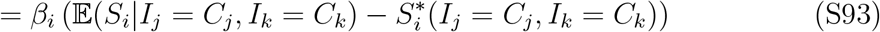

Therefore, a negative higher order term implies that the true amount of susceptible host density in the presence of two competitors is higher than the expected level under the linear assumption. In other words, the Lotka-Volterra model implicitly underestimates the amount of susceptible hosts for a newly invading strain.

A special case when strains *j* and *k* do not interact (thus, *Q*_*jk*_ = *Q*_*kj*_ = 0) presents an interesting example. In this case, both strains *j* and *k* will be able to coexist at their single strain equilibrium: 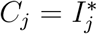 and 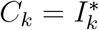. In this case, the differences in expected and realized susceptible host density corresponds to:

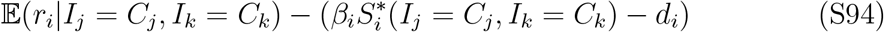

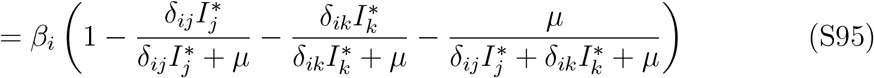

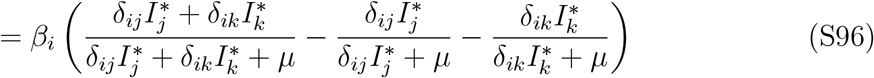

Rewriting 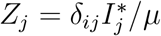 and 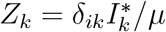, we have:

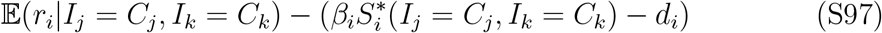

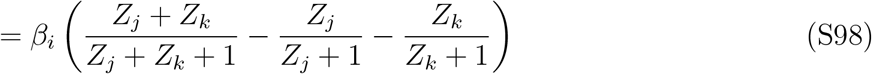

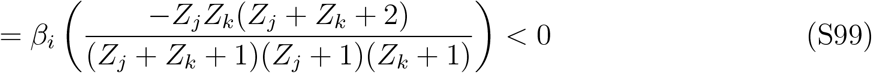

Therefore, even in the absence of interaction between strains *j* and *k* the higher order term will be always negative.

### S8 Deterministic simulation of rotavirus strain dynamics

We test the resulting framework against a realistic competitive system, using a classical model describing the competition of rotavirus strains, proposed by Pitzer et al. (2011). Briefly, the model describes the competition between 5 rotavirus strains, accounting for reduction in susceptibility and transmissibility with subsequent reinfections. Equations describing the underlying strain dynamics can be written as follows:

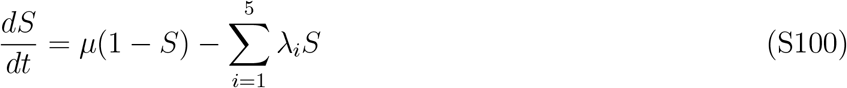

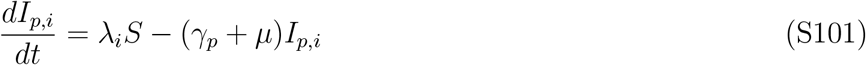

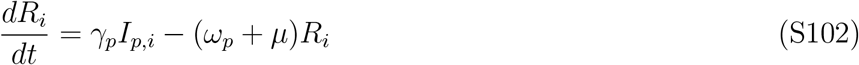

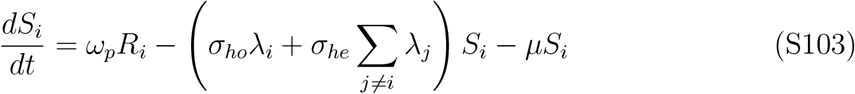

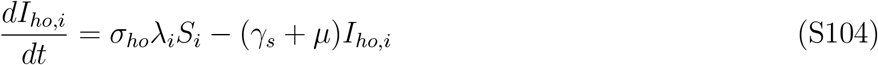

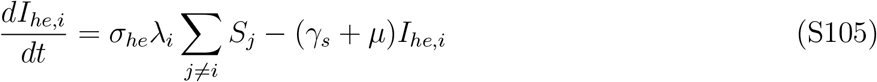

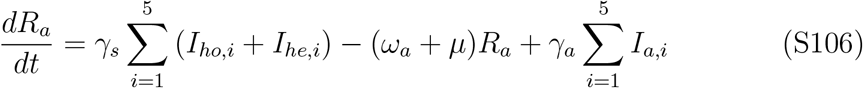

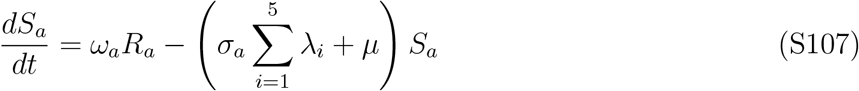

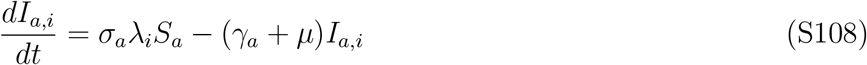

where *S* represents the proportion of susceptible individuals; *I*_*p,i*_ represents the proportion of primary infections with strain *i*; *R*_*i*_ represents the proportion of individuals who have recovered from a primary infection with strain *i* and are immune; *S*_*i*_ represents the proportion of individuals who have immunity against strain *i* and are susceptible to reinfections; *I*_*ho,i*_ represents the proportion of individuals who have been reinfected with strain *i* with immunity against the same strain; *I*_*he,i*_ represents the proportion of individuals who have been reinfected with strain *i* with immunity against a different strain; *R*_*a*_ represents the proportion of individuals who have recovered from a secondary infection and are immune; *S*_*a*_ represents the proportion of individuals who have recovered from a secondary infection and are susceptible to new infections; and *I*_*a,i*_ represents the proportion of asymptomatic infections with strain *i*. Then, *µ* represents the birth and death rates; *λ*_*i*_ represents the force of infection for strain *i*; *γ*_*x*_ represents recovery rates for infection class *x*; *ω*_*x*_ represents to immune waning rate for class *x*; *σ*_*ho*_ represents relative susceptibility to homotypic reinfection; *σ*_*he*_ represents relative susceptibility to heterotypic reinfection; and *σ*_*s*_ represents relative susceptibility after secondary infections. Here, the force of infection is modeled as follows:

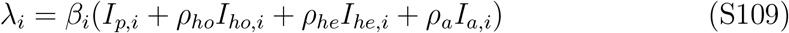

where *β*_*i*_ represents the transmission rate; *ρ*_*ho*_ represents relative transmissibility for homotypic reinfections; *ρ*_*he*_ represents relative transmissibility for heterotypic reinfections; and *ρ*_*a*_ represents relative transmissibility for tertiary reinfections. The original parameterization by Pitzer et al. (2011) assumed seasonally forced transmission to capture complex multiannual turnovers exhibited by different strains. Since our goal is to test whether a Lotka-Volterra model can accurately predict the equilibrium densities of a realistic competitive system, we neglect seasonal forcing for simplicity. Moreover, the original parameterization assumed high transmission for the first strain and equal, but lower transmission for all other strains; instead, we allowed different transmission rates across all strains so that each strain can have distinct equilibrium densities. Thus, we assume *beta*_1_ = 34*/*weeks, *beta*_2_ = 34*/*weeks × 0.98, *beta*_3_ = 34*/*weeks × 0.96, *beta*_4_ = 34*/*weeks × 0.94, and *beta*_5_ = 34*/*weeks × 0.92. For all other parameters, we assume 1*/ω*_*p*_ = 39 weeks, 1*/ω*_*a*_ = 52 weeks, 1*/γ*_*p*_ = 1 week, 1*/γ*_*s*_ = 0.5 weeks, 1*/γ*_*a*_ = 0.5 weeks, 1*/µ* = 50 × 52 weeks, *σ*_*ho*_ = 0.1, *σ*_*he*_ = 0.7, *σ*_*a*_ = 0.35, *ρ*_*ho*_ = 0.1, *ρ*_*he*_ = 0.7, and *ρ*_*a*_ = 0.1. We note that the system still exhibits a cyclic behavior even in the absence of seasonal forcing.

As a comparison, we also consider a model assuming polarized immunity. We model polarized immunity by including the immune boosting of exposed, but uninfected, individuals (Park et al., 2023). This is modeled assuming that all susceptible individuals experience identical forces of infections *λ*_*i*_, but individuals with partial susceptibility can either become infected with probability *σ*_*x*_ and gain full protection with probability 1 − *σ*_*x*_. As discussed earlier (Park et al., 2023), this formulation is equivalent to polarized immunity because the model implicitly assumes that a proportion 1 − *σ*_*x*_ of individuals with waned immunity are essentially immune to transmissible infections whereas the remaining proportion *σ*_*x*_ of individuals are fully susceptible to transmissible infections. Then, equations describing the underlying strain dynamics can be written as follows:

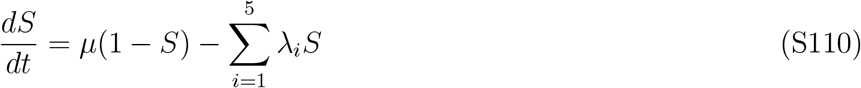

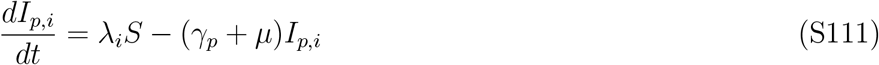

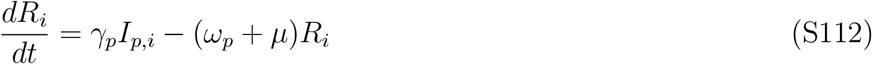

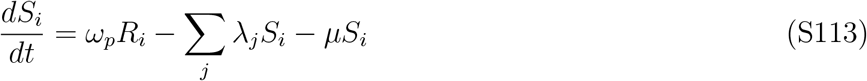

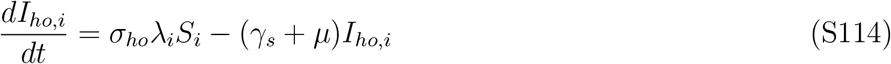

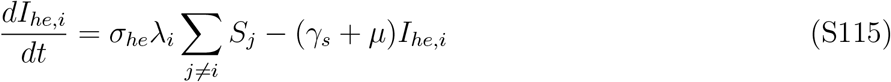

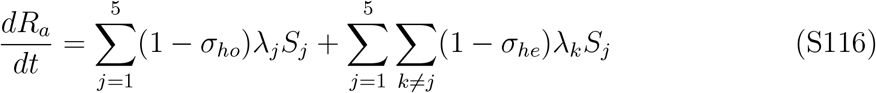

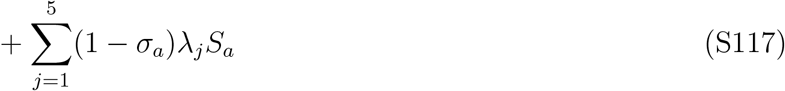

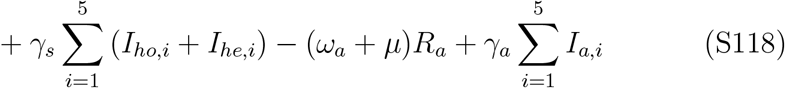

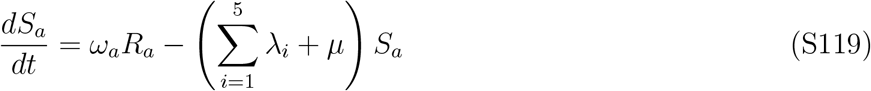

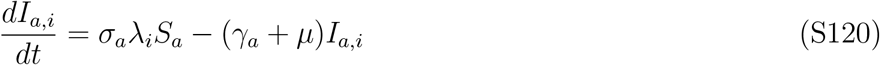

Note that the susceptible depletion always occurs at the per capita rate of 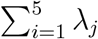 but some individuals develop transmissible infection, whereas others simply move to corresponding recovered compartments *R*. Parameters for this model are same as before.

Since both these models exhibit cyclic behavior, calculating the invasion growth rate is challenging. Thus, to remove the cyclic behavior, we speed up the dynamics of susceptible and recovered populations, which in turn limits susceptible resource overcompensation. Specifically, we multiply *dR/dt* and *dS/dt* by *s* = 10, which represents the relative speed of susceptible and recovered dynamics in comparison to infected dynamics. This approach is analogous to time scale separation used in earlier sections and is applied to both leaky and polarized immunity models.

By speeding up susceptible and recovered dynamics, we can now calculate the Lotka Volterra coefficients following the approach outlined in Section S4. In doing so, we use the proportion of primary infected individuals as the focal population density. We first calculate the equilibrium primary infections 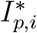 for each strain in the absence of competing strains and then introduce a competing strain *j*. Then, interspecific competition coefficients correspond to:

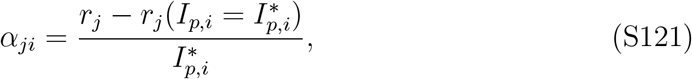

where *r*_*j*_ = *β*_*j*_ − (*γ*_*p*_ + *µ*) represents the intrinsic growth rate of strain *j* and 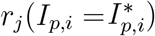 represents the growth rate of strain *j* when strain *i* is at equilibrium. Similarly, intraspecific competition coefficients correspond to:

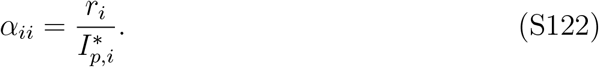

Then, we calculate higher order interaction terms by first allowing strains *j* and *k* to reach coexisting endemic equilibrium (*Î*_*p,j*_ and *Î*_*p,k*_), and introducing strain *i*:

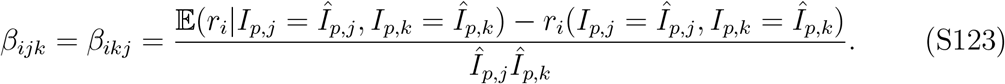

Here, 𝔼(*r*_*i*_|*I*_*p,j*_ = *Î*_*p,j*_, _*p,k*_ = *Î*_*p,k*_) represents the expected invasion growth rate of strain *i* when strains *j* and *k* are at equilibrium:

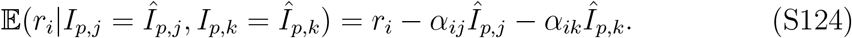

Similarly, *r*_*i*_(*I*_*p,j*_ = *Î*_*p,j*_, *I*_*p,k*_ = *Î*_*p,k*_) represents the realized growth rate of strain *i* when when strains *j* and *k* are at equilibrium—this is calculated by simulating the invasion of strain *i*. Note that we use the hat notation here to distinguish these coexisting equilibrium from single strain equilibrium. The intrinsic growth rate *r*_*i*_, pairwise interaction terms *α*_*ij*_ and higher order interaction terms *β*_*ijk*_ give full parameters to simulate the Lotka-Volterra model. In the actual Lotka-Volterra simulations, we divided all three terms by 10 to avoid integration error due to large parameters by slowing down the system—since we are only interested in equilibrium densities, these time scale slowing down does not affect our conclusion.

## Notes

### Competing Interest Statement

The authors have declared no competing interest.

